# Integrating continuous hypermutation with high-throughput screening for optimization of *cis,cis*-muconic acid production in yeast

**DOI:** 10.1101/2020.12.09.418236

**Authors:** Emil D. Jensen, Francesca Ambri, Marie B. Bendtsen, Alex A. Javanpour, Chang C. Liu, Michael K. Jensen, Jay D. Keasling

## Abstract

Directed evolution is a powerful method to optimize proteins and metabolic reactions towards user-defined goals. It usually involves subjecting genes or pathways to iterative rounds of mutagenesis, selection, and amplification. While powerful, systematic searches through large sequence-spaces is a labor-intensive task, and can be further limited by *a priori* knowledge about the optimal initial search space, and/or limits in terms of screening throughput. Here we demonstrate an integrated directed evolution workflow for metabolic pathway enzymes that continuously generates enzyme variants using the recently developed orthogonal replication system, OrthoRep, and screens for optimal performance in high-throughput using a transcription factor-based biosensor. We demonstrate the strengths of this workflow by evolving a rate-limiting enzymatic reaction of the biosynthetic pathway for *cis*,*cis*-muconic acid (CCM), a precursor used for bioplastic and coatings, in *Saccharomyces cerevisiae*. After two weeks of simply iterating between passaging of cells to generate variant enzymes via OrthoRep and high-throughput sorting of best-performing variants using a transcription factor-based biosensor for CCM, we ultimately identified variant enzymes improving CCM titers >13-fold compared to reference enzymes. Taken together, the combination of synthetic biology tools as adopted in this study, is an efficient approach to debottleneck repetitive workflows associated with directed evolution of metabolic enzymes.

## Introduction

Industrial biotechnology has offered commercialization of environmentally friendly transportation fuels, amino acids and value-added chemicals by the use of fermentation feedstocks and microbial cell factories (Choi *et al*., 2019). Yet, industrializing microbial cells for a broad range of applications within manufacturing, health, and transportation industries often requires extensive engineering of both the microbial chassis as well as at the level of scaling up the fermentation processes (Van Dien, 2013; Nielsen and Keasling, 2016). Indeed, in the design of cell factories for fermentation-based manufacturing of value-added chemicals and therapeutics, biosynthetic pathways are often composed of enzymes from several different sources, and with enzyme activities and expression levels requiring careful balancing in order to achieve optimal pathway flux (Galanie *et al*., 2015; Zhang *et al*., 2020). While such multi-dimensional optimization can be streamlined using design-of-experiment approaches and machine learning algorithms (Jeschek *et al*., 2016; Xu *et al*., 2017; Carbonell *et al*., 2018), the regulatory and cellular complexity of living cells and the constraints in speed, scale, depth, and costs of even rational trial-and-error engineering approaches challenge the development of microbial cell factories.

As a complementary approach to bottom-up rational engineering, evolution-guided cell factory engineering has gained substantial traction over the last decade (Mundhada *et al*., 2016; Sandberg *et al*., 2019). Here, the key principles of evolutionary engineering includes targeted or genome-wide genetic diversification coupled with screening of variant libraries (Packer and Liu, 2015). Numerous metabolic engineering studies have successfully applied directed evolution to improve product and feedstock tolerance as well as cell factory performance (Caspeta *et al*., 2014; Park *et al*., 2014; Mundhada *et al*., 2016). While powerful, both the generation of large numbers of genetic variants and the development of proper selection regimes, as well as the cloning and transformation procedures associated with directed evolution cycles, are often time- and cost-intensive. To overcome this, *in vivo* directed evolution uses endogenous or orthogonal cellular machineries to maintain high mutation rates without the need for iterative cycles of library cloning and transformation, apart from propagating the evolving population (Esvelt *et al*., 2011; Ravikumar *et al*., 2014; Crook *et al*., 2016). One such system is OrthoRep enabling continuous generation of variant genes of interest expressed from a linear cytoplasmic chromosome that is propagated via an orthogonal error-prone DNA polymerase (Ravikumar *et al*., 2014, 2018). With orthogonal *in vivo* evolution machineries at hand, any trait that can be coupled to growth (e.g. antibiotic resistance, tolerance to cultivation conditions, and/or complementation of auxotrophies) enables facile identification of improved target genes without need for direct screening (Esvelt *et al*., 2011; Ravikumar *et al*., 2014; García-García *et al*., 2020; Rix *et al*., 2020).

However, for metabolic engineering, the expression of heterologous enzymes and proteins towards biobased production of value-added chemicals seldom allows direct coupling of production to growth, or other high-throughput screens, needed to capitalize on the massive diversity generated by *in vivo* evolution systems (Esvelt and Wang, 2013). Here the recent development of biosensors based on allosterically regulated transcription factors (aTFs) can provide a complementary technology for coupling enzymatic activity or pathway flux with facile screening of large variant libraries in multiplex through fluorescence-activated cell sorting (FACS) or growth (Raman *et al*., 2014; Flachbart *et al*., 2019). Briefly, such biosensors link binding of small molecule ligands to aTFs as input, with changes in expression of reporter genes or actuators as output (Mahr and Frunzke, 2016). Taken together, the coupling of continuous evolution systems with biosensing could allow metabolic engineers to cost-effectively search for optimal pathway designs.

Here we combine the power of targeted *in vivo* mutagenesis using OrthoRep with high-throughput biosensing for the rapid evolution of rate-limiting metabolic reactions of the *cis,cis-muconic* acid (CCM) pathway (Weber *et al*., 2012; Curran *et al*., 2013; Suástegui and Shao, 2016). The 3-step CCM pathway, consisting a dehydroshikimate dehydratase (AroZ), a multi-subunit protocatechuic acid (PCA) decarboxylase, and a catechol 1,2 dioxygenase (**Figure 1A**) (Weber *et al*., 2012; Curran *et al*., 2013), has been extensively studied and optimized to support biobased production of plastics and coatings, following hydrogenation of CCM into adipic acid as a building block for nylon-6,6 (Weber *et al*., 2012; Curran *et al*., 2013; Suastegui *et al*., 2016; Leavitt *et al*., 2017; Snoek *et al*., 2018; Wang *et al*., 2020). Importantly, for CCM pathway optimization, we have previously engineered the aTF BenM as a CCM biosensor in yeast (Skjoedt *et al*., 2016), and this has further enabled optimization of yeast as a chassis for CCM production (Snoek *et al*., 2018; Wang *et al*., 2020), complemented by additional evolution- and machine learning-guided optimization of the endogenous yeast aromatic amino acid pathway from which CCM is derived (Leavitt *et al*., 2017; Zhang *et al*., 2020). However, the build-up and secretion of the CCM pathway intermediate PCA remains rate-limiting for high CCM production (Suastegui *et al*., 2016; Leavitt *et al*., 2017; Snoek *et al*., 2018; Wang *et al*., 2020). This observation is propelled by the fact that PCA accumulation supposedly is not limited to suboptimal catalytic activity or expression of the downstream CCM pathway enzyme, PCA decarboxylase, as PCA accumulation also remains a persistent issue for microbial biosynthesis of other PCA-derived chemicals and nutraceuticals without decarboxylase requirements (Strucko *et al*., 2015, 2017; D’Ambrosio *et al*., 2020). Yet, no attempts for directed evolution of the heterologous enzymes towards increased PCA-derived production has to our knowledge been performed. Here we demonstrate rapid evolution of PCA decarboxylase subunits using a simple experimental design enabled by OrthoRep and five biosensor-assisted selection cycles ultimately yielding >13-fold higher CCM production compared to the subunits encoded in the wild-type PCA decarboxylase complex.

**Figure 1.**
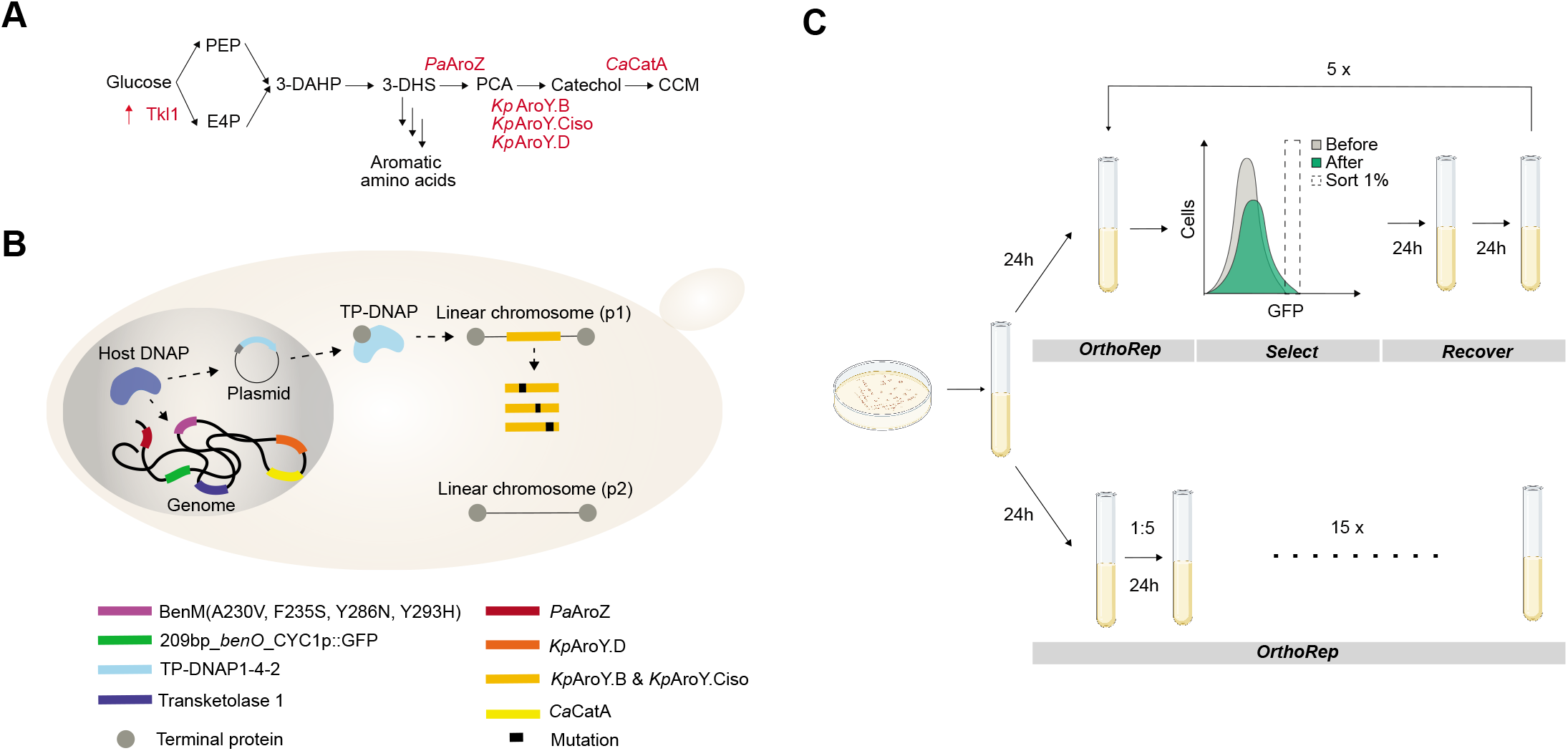
Schematic illustration of the *n vivo* directed evolution workflow. (**A**) Schematic illustration of the 3-step *cis*,*cis*-muconic acid pathway, comprising heterologous expression of *PaAroZ*, *KpAroY* subunits (B, D, and Ciso), as well as *CaCatA* and overexpression of Tkl1 (Weber *et al*., 2012; Curran *et al*., 2013). (**B**) Schematic illustration of the parental strain (Sc-105, see **Supplementary Table S5**) used for *in vivo* directed evolution of the *cis*,*cis*-muconic acid pathway enzymes *KpAroY.B* and *KpAroY.Ciso* in yeast cells. The strain replicates and expresses the biosensor, all *cis*,*cis*-muconic acid pathway enzymes except *KpAroY.B* and *KpAroY.Ciso*, and the variant error-prone TP-DNAP (expressed from AR-Ec633, see **Supplementary Table S4**) from the nucleus. All components required for OrthoRep replication and transcription are encoded on p2, whereas genes encoding *KpAroY.B* and *KpAroY.Ciso* are expressed from p1. (**C**) Schematic illustration of the *in vivo* directed evolution workflow showing the passaging regimes of the parental strain undergoing i) the five consecutive rounds of OrthoRep coupled with biosensor-based selection, or ii) fifteen bulk passages to effect drift without biosensor-based selection.

## Results

### Parental strain design

In order to efficiently evolve *cis*,*cis*-muconic acid biosynthetic pathway enzymes we used state-of-the-art orthogonal error-prone replication and biosensing machineries. For generation of sequence diversity using OrthoRep, we used the highly error-prone TP-DNAP1 variant, TP-DNAP1(L477V, L640Y, I777K, W814N), with mutation rate of ~1×10^-5^ substitutions per base (Ravikumar *et al*., 2018), whereas for selection of high-performing *cis*,*cis*-muconic acid pathway designs we used the CCM-binding BenM variant transcription factor, BenM(A230V, F253S, Y286N, Y293H), with a high dynamic output range to maximise sorting resolution (Snoek *et al*., 2019). While the TP-DNAP1 variant was expressed from a CEN/ARS-based plasmid, the BenM variant was genomically integrated together with an expression cassette encoding the GFP reporter (**Figure 1A**) (Ravikumar *et al*., 2018; Snoek *et al*., 2018). With respect to the CCM pathway template, we genomically integrated codon-optimized *AroZ* from *Podospora anserina (PaAroZ), CatA* from *Candida albicans* (*CaCatA*), and the gene encoding the D subunit of *AroY* from *Klebsiella pneumoniae* (*KpAroD*) for heterologous expression of the 3-DHS dehydratase, catechol 1,2-dioxygenase, and D subunit-mediated PCA decarboxylase reactions, respectively (Weber *et al*., 2012; Curran *et al*., 2013)(**Figure 1A**). In order to evolve the rate-limiting PCA decarboxylase reaction using OrthoRep, we encoded the codon-optimized B subunit and the C isoform subunit without oxygen sensitivity of *AroY* from *K. pneumoniae (KpAroY.B* and *KpAroY.Ciso*, respectively) (Weber *et al*., 2012) on the p1 linear plasmid for replication with the error-prone TP-DNAP1 variant. Together with overexpression of the transketolase gene *(TKL1)* to increase the carbon flux into the aromatic amino acid biosynthesis (Luttik *et al*., 2008; Curran *et al*., 2013), this parental strain (Sc-105) was used as a starting point for *in vivo* directed evolution of CCM production in yeast (**Figure 1B**).

### Selection of hypermutated CCM biosynthetic pathway enzymes

In order to evolve the parental strain towards higher CCM production, we coupled continuous mutagenesis using OrthoRep with selection using the BenM-based biosensor output as a proxy for CCM accumulation, respectively. Specifically, with the parental starting strain Sc-105 in place, we cultured cells in Delft minimal medium to allow OrthoRep to generate AroY diversity and selected high CCM-producing cells by fluorescence-activated cell sorting (FACS) of 1 mio. events. We iterated culturing and FACS five times, where each sort collected 10,000 events and was followed by a 24-hr recovery cultivation in synthetic complete medium, after which propagated cells were harvested for both glycerol stock and passed for further evolution. For this conservative selection schedule, each such cycle required 3 days. As a control evolution experiment, we cultured the parental starting strain over 15 consecutive 24 hrs cultivations without FACS selection by bulk passaging 20% of cells between each cultivation. By monitoring population-level fluorescence changes in this control evolution experiment, we could ensure that no changes in CCM production resulted from simply drifting AroY on OrthoRep. In total, both regimes were conducted over 15 days, totaling 5 sorted populations and 15 bulk populations (**Figure 1C**).

The number and duration of the iterative cycles were selected based on i) the mutation-rate and copy number of TP-DNAP1-4-2, ii) the sequence length of the bait genes encoding AroY.B and AroY.Ciso, and iii) the generation time of *S. cerevisiae* (Ravikumar *et al*., 2018). Briefly, with an estimated mutation-rate of 1 x 10^-5^ substitutions per base at 10x copies of p1 per cell (Ravikumar *et al*., 2018), the actual mutation-rate per cell approximates 1 x 10^-4^ substitutions per base. With *AroY.B* and *AroY.Ciso* sequences on the p1 plasmid totalling 2.1 kb (**Supplementary Table S1, Supplementary Figure S1**), we expected 0.21 mutations/replication/generation, or 1 mutation every fifth generation. In addition to this, for the actual cultivations, we assumed that no mutations would occur during the first 24 hrs recovery following each sort, and that the exponentially growing cells would double every ~3 hrs reaching a max OD600 (optical density measured at a wavelength of 600 nm) of 8 following 5-6 doublings in Delft minimal medium before each sort. These experimental choices would on average result in approximately 1 additional mutation accumulated in *AroY.B* and *AroY.Ciso* per cell for each additional sort.

### Characteristics and validation of selected enzyme variants

Following the five iterations of OrthoRep coupled with selection using FACS, we compared the population-level fluorescence distributions from the five sorted populations as well as the 15 bulk passaged populations (**Figure 2A**). Over the course of the evolution experiment the population-level fluorescence outputs increased following each round of sorting from the first to the fourth sort, reaching a total increase of 2.5-fold. From the fourth to fifth sorting no further increase in fluorescence was observed (**Figure 2A**). In total, a maximum of 3.9-fold increase was observed when comparing mean fluorescence outputs following five rounds of sorting versus fifteen rounds of neutral drifting (**Figure 2A**, “Sort 5” vs “Bulk 15”). With the interest to inspect fluorescence outputs of cells undergoing drifting vs selection, we isolated 20 colonies obtained from “Bulk 15” and >300 colonies obtained from “Sort 4+5”, respectively. Here, the increased fluorescence was confirmed, with an average fluorescence output of colonies from sorted populations >5-fold higher than the average fluorescence output of colonies from cells undergoing neutral drifting (**Figure 2B**, insert). Importantly, and corroborating the conservative selection regime, more than 96% (309/319) of the colonies from sorted populations (“Sort4+5”) had cells with higher fluorescence than cells from colonies undergoing neutral drifting (“Bulk 15”) (**Figure 2B**, dashed line).

**Figure 2.**
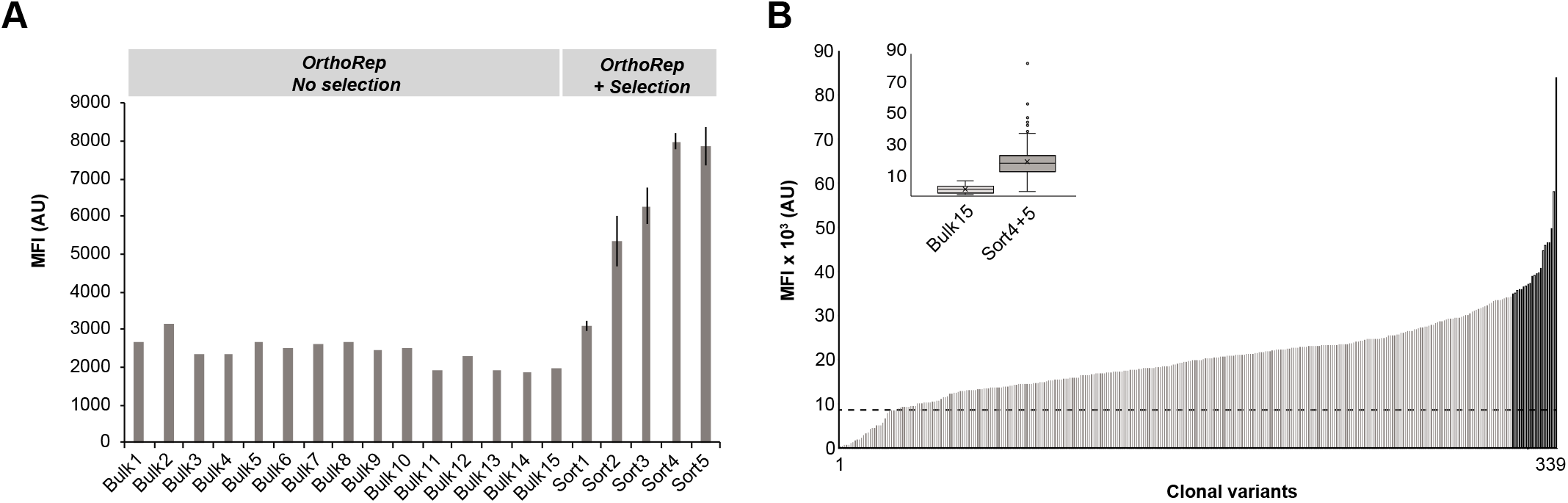
Population-level fluorescence outputs from parental strains expressing evolvable PCA decarboxylase subunits. (**A**) Population-level fluorescence outputs following 15 passages of cultures of parental strains undergoing neutral drifting (Bulk 1-15) by OrthoRep (OrthoRep - No selection), and following 5 consecutive iterations (Sort 1-5) of OrthoRep coupled to CCM biosensor-based selection (OrthoRep + Selection). Bars indicate mean fluorescence intensity (MFI) of 10,000 events. AU: arbitrary units. (**B**) Mean fluorescence intensity of cells from 20 colonies propagated from population “Bulk 15” and 309 colonies propagated from Sort 4 and Sort 5 (“Sort4-5”). Insert shows box plots from all 20 and 309 mean fluorescence intensities obtained from the 10,000 events measured for each of the “Bulk15” and “Sort4+5” populations, respectively. Bars indicate mean fluorescence intensity (MFI) of 10,000 events. Error bars represent standard deviation of the mean from three biological replicate samplings. AU: arbitrary units.

Next, from the >300 colonies re-screened using flow cytometry (**Figure 2B**), we sequenced the *KpAroY.B* and *KpAroY.Ciso* alleles amplified from the p1 plasmid of cells derived from 30 of the highest fluorescent colonies (**Figure 2B**, dark bars). Here we found 11 amino acid changes, 2 deletion, and 9 silent mutations (**Figure 3A**), with all mutations occurring only once, except the *KpAroY.Ciso_666T>del* (deletion of single nucleotide following 666T, leading to stop codon at residue 244), occurring twice. Of the missense and nonsense mutations, four were located in *KpAroY.B* and nine in *KpAroY.Ciso* (**Figure 3A, Supplementary Table S2**). To validate that the evolved *KpAroY.B* and *KpAroY.Ciso* mutant alleles, and not other potential mutations in the p1 plasmid or nuclear genome, were causal for the observed increase in fluorescence, we cloned individual *KpAroY.B* and *KpAroY.Ciso* alleles into the genome of a parental strain (Sc-78) under the control of the strong constitutive promoters TDH3 and TEF1, respectively (**Supplementary Figure S2**). For each *KpAroY.B* variant allele, a wild-type *KpAroY.Ciso* allele was included in the integration, and *vice-versa*. The parental strain harboured the *PaAroZ, CaCatA* and CCM biosensor genes genomically integrated, and fluorescence and metabolites compared with the parental strain expressing wild-type *KpAroY.B* and *KpAroY.Ciso*. A total of 8 *KpAroY* variants, 2 *KpAroY.B* and 6 *KpAroY.Ciso*, were tested and compared with the strain expressing wild-type *KpAroY.B* and *KpAroY.Ciso*. Firstly, we tested fluorescence of the 8+1 strains following 24 hrs of cultivation. Here we found that only 5 of the 8 variants showed significantly higher fluorescence outputs compared to the wild-type *KpAroY* reference strain (p-value < 0.01), with *KpAroY.B_P146T* performing the best (10.7x) (**Figure 3B**). Next, we analysed the amounts of CCM and PCA of all strains following 72 hrs of cultivation, as previously shown to be a relevant benchmark time-point for comparing biosensor fluorescence outputs and CCM titers (Skjoedt *et al*., 2016). Here, we observed that 6 out of 8 strains showed significantly higher CCM titers compared to the wild-type *KpAroY* reference strain (*t*-test, p-value < 0.01), again with *KpAroY.B_P* 146T performing the best with a 13.7-fold higher CCM titer when expressed together with wild-type *KpAroY.Ciso*, compared to expressing wild-type *KpAroY.B* and *KpAroY.Ciso* together (**Figure 3C**). Interestingly, when comparing fluorescence and CCM titers, it becomes clear that while the CCM biosensor variant used in this study (BenM(A230V, F253S, Y286N, Y293H)) indeed enables selection of *KpAroY.B* and *KpAroY.Ciso* variant alleles yielding higher CCM production, the biosensor operational range cannot discriminate CCM titers higher than approximately 80 mg/L under this cultivation regime (**Figure 3B-C**). While this is an inherent limitation of the specific BenM variant, it is critical to underscore that this variant was chosen due to its high dynamic output range (Snoek *et al*., 2019), which was deemed necessary for selection in multiplex. Lastly, having verified that CCM product titers were increased when expressing 6 of the 8 *KpAroY* variants, we next investigated if PCA titers for the pathways expressing PCA decarboxylase variants were perturbed. Importantly, PCA titers of no higher than 250-300 mg/L are known to cause a significant fitness burden to *S. cerevisiae (D’Ambrosio et al., 2020)*, making it critical to investigate if the observed increases in CCM titers for the 6 different PCA decarboxylase variants would support catalytic activities to lower PCA below fitness-burdening PCA levels. Indeed, of the *KpAroY* variants supporting increased CCM titers, *KpAroY.B*_P146T also showed significantly reduced PCA titers (77%, p < 0.01)(**Figure 3D**), further highlighting this variant as a new and catalytically improved PCA decarboxylase.

**Figure 3.**
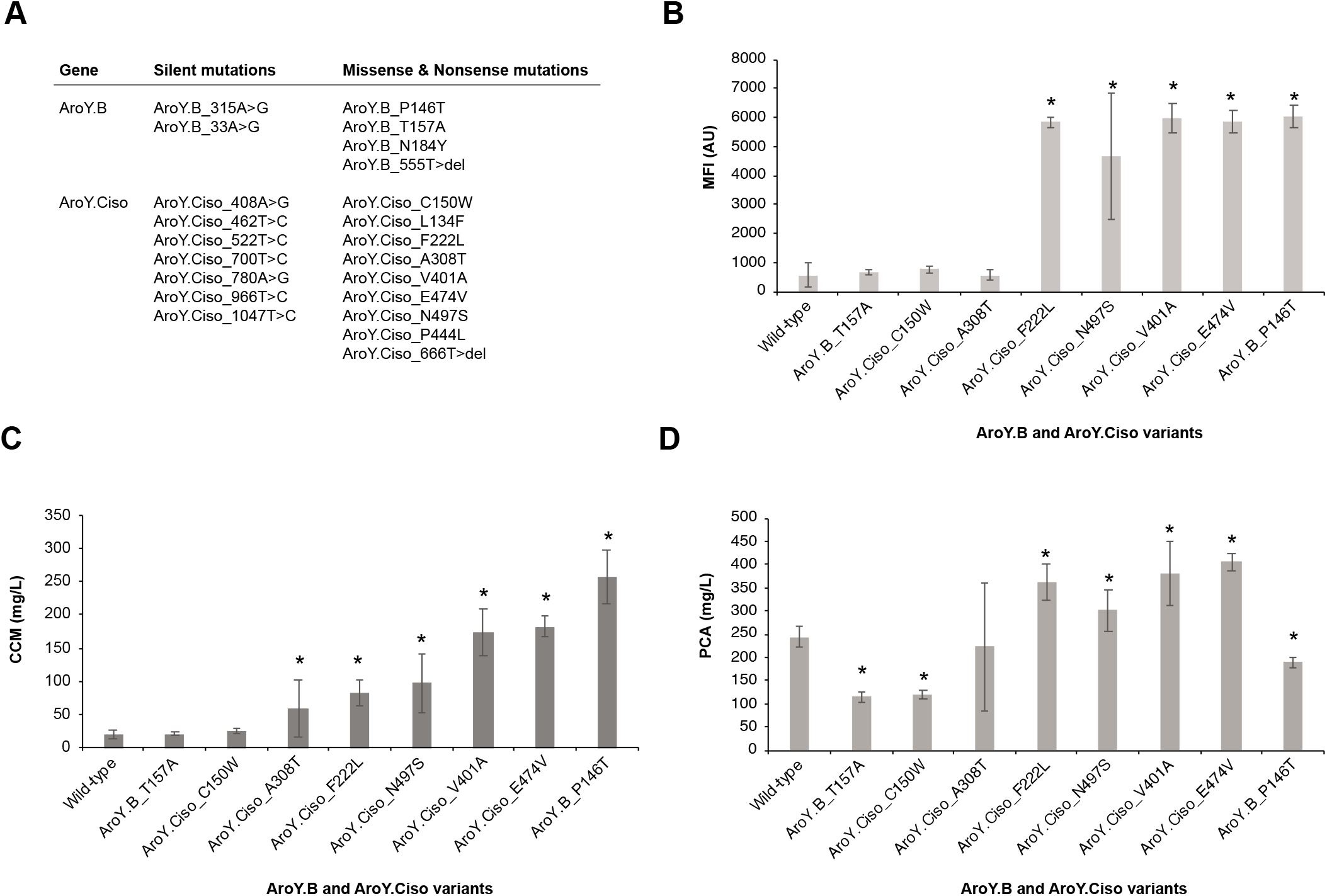
Characterization of PCA decarboxylase subunit variants from OrthoRep-evolved populations. (**A**) Parent populations from which OrthoRep-evolved *KpAroY.B* and *KpAroY.Ciso* variants derived, and the silent, missense and nonsense mutations present in them. (**B**) CCM biosensor-based evaluation of 8 evolved PCA decarboxylase subunit variants *KpAroY.B* and *KpAroY.Ciso* compared to wild-type PCA decarboxylase subunits through flow cytometry assays. Mean fluorescence intensity (MFI) following 24 hrs cultivation for parental strain Sc-78 integrated with the indicated 8 evolved *KpAroY.B* and *KpAroY.Ciso* PCA decarboxylase subunit variants or wild-type PCA decarboxylase subunits (Sc-194). Data represent means of 4-5 biological replicates, and error bars represent standard deviation of the mean. (**C**) Extracellular *cis,cis*-mucinic acid (CCM) concentrations in cultivation broth from the same as in (B) 8 evolved PCA decarboxylase subunit variants of *KpAroY.B* and *KpAroY.Ciso* compared to wild-type PCA decarboxylase subunits following 72 hrs cultivation. Data represent means of 4-5 biological replicates, and error bars represent standard deviation of the mean (**D**) Extracellular protocatechuic acid (PCA) concentrations in cultivation broth from the same as in (B) and (C) 8 evolved PCA decarboxylase subunit variants *KpAroY.B* and *KpAroY.Ciso* compared to wild-type PCA decarboxylase subunits following 72 hrs of cultivation. Data represent means of 4-5 biological replicates, and error bars represent standard deviation of the mean.

## Discussion

In this study we demonstrate the successful merger of two synthetic biology tools for the benefit of evolving superior enzymes without the use of labor-intensive library designs or costly low/semi-throughput analytical facilities. Importantly, this study show-cases the evolution of metabolic pathway enzymes without any native growth advantage for the cells, a condition that most previous *in vivo* directed evolution requires (Esvelt *et al*., 2011). Furthermore, the merger of OrthoRep and biosensors for directed evolution as demonstrated in this study complements the development of selections associated with growth under (strong) selection pressures (Ravikumar *et al*., 2018; Zhong *et al*., 2020).

While this study was a successful demonstration of continuous hypermutation of target genes applied for metabolic enzyme evolution, more prospecting and evolution-guided engineering of the CCM pathway is still warranted. For biobased CCM production, such efforts should not be limited to improving the suboptimal catalytic activity or expression of pathway enzymes downstream of PCA, i.e. PCA decarboxylase and catechol 1,2-dioxygenase. One strategy to consider further is the need to limit the secretion, or passive diffusion, of PCA across the cellular membrane as is often observed in yeast engineered to produce PCA-derived chemicals and nutraceuticals (Hansen *et al*., 2009; Weber *et al*., 2012; Curran *et al*., 2013; Suastegui *et al*., 2016; Leavitt *et al*., 2017). With the evolved PCA decarboxylase subunits identified from this study, a logical next step could be to move from *in vivo* directed evolution of predefined target genes towards genome-wide adaptive laboratory evolution. For such purposes, transferring the biosensor read-out from fluorescence to growth would be beneficial from a technical and scalability point of view (Zhong *et al*., 2020). For such a purpose, the coupling of CCM to growth using synthetic control circuits founded on BenM, or the recently developed vanillin biosensor, could be useful for genome-wide searches of nucleotide polymorphisms and chromosomal rearrangements limiting PCA efflux and/or further boosting PCA metabolic flux, respectively (Ambri *et al*., 2020; D’Ambrosio *et al*., 2020).

Another aspect to consider is to diversify the selection criteria beyond the stringent one used in this study (**Figure 1C**). Specifically, tuning the selection strength during continuous evolution regimes has previously been demonstrated to enable mutational drifting and adaptation of robust proteins (Bershtein *et al*., 2008; Steinberg and Ostermeier, 2016; Zhong *et al*., 2020). With the coupling of OrthoRep to FACS-compatible screens as demonstrated in this study, such tuning should be possible to implement, and further explored, in order not to outpace the rate of adaption using the conservative and stringent cut-off for selection as applied in this study. Ultimately, this could expand both the robustness and catalytic activity of further evolved PCA decarboxylases, but also increase the relatively low number of mutations observed per evolved PCA decarboxylase variant. Furthermore, toggled selection regimes of neutral drifting interrupted by selection (Rix *et al*., 2020; Zhong *et al*., 2020), may also increase the hit-rate of the continuous evolution, and limit the false-discovery rate observed in this study (>9/22, >0.45)(**Figure 3**).

In summary, we consider that our study serves as a first demonstration of rapid evolution of metabolic enzymes without any direct fitness advantage using continuous hypermutation, and furthermore moves forward the engineering of *S. cerevisiae*, and potentially other microbial chassis, for the industrial production of CCM as a precursor for further hydrogenation into adipic acid and nylon-6,6 for the bioplastics industry.

## Experimental Procedures

### Cloning

Plasmid pEDJ366 carries a *URA3* targeting gRNA and was made by inverse amplification of pCfB3050 (Jessop-Fabre *et al*., 2016) with oligos EDJ382 and EDJ383 followed by T4 DNA ligation (NEB). Plasmid p1237 (Skjoedt *et al*., 2016) was modified for *SpHIS5::TRP1* by inverse amplification with oligos EDJ386 and EDJ387, and assembled with CEN.PK2-1C genomic *TRP1* amplified with EDJ388 and EDJ389 by T4 DNA ligation (pEDJ371). pCfB2764 was amplified with EDJ432 and EDJ433 and assembled with gEDJ12 obtained with primers EDJ434 and EDJ435 to give plasmid pEDJ515. gEZ475 was amplified by EDJ390 and EDJ391 and assembled with ZZ-Ec475 (Zhong *et al*., 2020) after restriction digest by *XhoI* and *XbaI* (pMB10). The p1 (Ravikumar *et al*., 2014) integration cassette (**Supplementary Table S1**) that contains wild-type *KpAroY.B* and *KpAroY.Ciso* was made by USER assembly (Jensen *et al*., 2014) with pMB10 as backbone vector and *KpAroY.B* and *KpAroY.Ciso* amplified from p1241 (Skjoedt *et al*., 2016) with primer pairs EDJ413 and EDJ414, and EDJ415 and EDJ416, respectively (pMB11). All oligonucleotides used in this study can be found in **Supplementary Table S3**.

### Media

One L of mineral medium (Delft) with 2% glucose (Verduyn *et al*., 1992) contained 75 ml (NH_4_)_2_SO_4_ (100 g/L), 120 ml KH_2_PO_4_ (120 g/L), 10 ml MgSO_4_, 7H_2_O (50 g/L), 2 ml trace metals, 1 ml vitamins, and 20 g glucose. 1 L of trace metals contain 4.5 g CaCl_2_·2H_2_O, 4.5 g ZnSO_4_·7H_2_O, 3 g FeSO_4_·7H_2_O, 1 g H_3_BO_3_, 1 g MnCl_2_·4H_2_O, 0.4 g Na_2_MoO_4_·2H_2_O, 0.3 g CoCl_2_·6H_2_O, 0.1 g CuSO_4_·5H_2_O, 0.1 g KI, and 15 g EDTA. 1 L of vitamins contain 50 mg biotin, 200 mg p-aminobenzoic acid, 1 g nicotinic acid, 1 g Ca-pantotenate, 1 g pyridoxine HCl, 1 g thiamine HCl, and 25 g myo-Inositol. Synthetic complete dropout media were bought from Sigma-Aldrich.

### Strains

Strain CEN.PK2-1C (EUROSCARF) expressing pRS414-TEF1p-Cas9-CYC1t (DiCarlo *et al*., 2013) was transformed with gRNA plasmid pEDJ366 and URA3-KO-90-mer to completely remove *ura3-52* (Sc-62). CCM-producing strain Sc-78 was made essentially as previously described (Skjoedt *et al*., 2016) with minor modifications; *NotI* treated plasmids p1237 and pEDJ371 were sequentially integrated in Sc-62 to give Sc-67 and Sc-68, followed by integration of similarly treated pCfB2553 and pEDJ515. Sc-78 was transformed with AR-Ec633 (Ravikumar *et al*., 2018) and saved as Sc-79 from histidine dropout medium. F102-2 was transformed with *Sca*I treated pMB11 (Sc-93), and resulting transformants were picked from uracil drop-out medium. Sc-93 was protoplast fused with Sc-79 as previously described (Ravikumar *et al*., 2014) and selected in synthetic complete histidine and uracil dropout medium to give Sc-105. Screened variants and wild-type *KpAroY.B* and *KpAroY.Ciso* (Sc-194) were synthesized as gene blocks by IDT and integrated with overlapping homology into Easyclone site XII-5 (Jensen *et al*., 2014) in strain Sc-78 with Cas9 and pCfB3050. Oligos ORP3-10 were used for amplifying individual parts from gene blocks or genomic DNA as indicated in **Supplementary Table S3** prior to reverse engineering of variant enzymes, and ORP1 with ORP2 and ORP11 with ORP12 for amplification of terminators with chr:XII-5 homology from pCfB2909 (Jessop-Fabre *et al*., 2016).

All plasmids and strains designed and constructed in this study can be found in **Supplementary Tables S4-S5**.

### Flow cytometer analysis

Strains were inoculated into appropriate synthetic complete dropout media (Sigma Aldrich) and incubated for 48 hrs in 96 deep-well plates at 30 °C with shaking. All cultures were then diluted 10-fold in fresh synthetic complete dropout media and incubated as before for 24 hrs or 72 hrs as indicated. All cultures were diluted 5-fold in a total volume of 150 μl with 1x phosphate buffer saline (PBS) from Life Technologies immediately before analysis. The BD LSRFortessa™ from BD Biosciences was used for analysing 10,000 single events per culture with a blue laser at 488 nm and settings FITC: 450, FSC: 150, and SSC: 250. FlowLogic (Invai Technologies) was used for processing of collected data.

### Mutation analysis

*KpAroY.B* and *KpAroY.Ciso* amplicons, derived with primers p1F and p1R from the p1 plasmid, were Sanger sequenced on individual clones. Three primers were used for sequencing to cover both genes (p1F, p1S, and p1R). The sequencing trace files were analyzed with non-evolved p1 plasmid as a reference in mutation surveyor using default settings (SoftGenetics) (Minton *et al*., 2011). Poor-quality sequences at the beginning and the end of the trace files were trimmed, and all called mutations were manually verified. All mutant sequences are listed in **Supplementary Table S2**.

### Fluorescence-assisted cell sorting

Parental strain Sc-105 was streaked on synthetic complete histidine and uracil dropout plates and incubated at 30 °C for 3 days; a single colony was grown in 5 mL of fresh Delft minimal medium for 24 hours at 30 °C with shaking. Pre-inoculo culture was obtained by diluting individual cell culture to OD of 0.2 in a total volume of 5 mL fresh Delft minimal medium and grown for an additional day at 30 °C with shaking. The culture was diluted again to OD of 2 in a total volume of 5 mL fresh Delft minimal medium and grown for another day at 30 °C with shaking. After 24 hours, 1 mL of the culture was named “Bulk 1” and saved as glycerol stock at −80°C; 1 mL was diluted in a total volume of 5 mL fresh Delft minimal medium to continue the “Bulk” cell culture and grown at 30 °C with shaking; and the remaining of the culture was diluted to OD of 0.5 in a total volume of 1 mL phosphate-buffered saline (PBS) in sterile tubes to arrest cell growth for flow cytometry acquisition. Using a SH800S cell sorter (Sony Biotechnology), 1 million events were analysed with a blue laser at 488 nm, and 10,000 cells of the top 2% fluorescent output were sorted into 2 mL of fresh synthetic complete medium and grown for 24 hours at 30 °C with shaking. The sorted population was then spun down and resuspended in 2 mL of fresh Delft minimal medium for additional 24 hours of recovery at 30 °C with shaking. Finally, 1 mL of the culture was named “Sort 1” and saved as glycerol stock at −80°C; the remaining population was diluted to OD of 2 in a total volume of 5 mL Delft minimal to be grown again for flow cytometry acquisition and additional sorting, totalling 5 “Sort” samples over the course of the 2 weeks experiment (**Figure 1B**). The “Bulk” culture was reincolutated daily into a total volume of 5 mL fresh Delft minimal medium totalling to 15 “Bulk” samples. Since the “Bulk” samples were not subjected to fluorescence enrichment they represent the neutrally drifted strains as a control for the sorted populations. The experiment has been done in triplicates, and 14 mL Falcon™ Round-Bottom Polypropylene Tubes (Thermo Fisher Scientific) were used for every cultivation.

### HPLC

Three replicates from each individual test strain were inoculated in 200 μL mineral medium (pH 4.5) supplemented with 20 mg/L Histidine and Uracil in 96 deep-well plates with air-penetrable lids (EnzyScreen) and incubated for 24 hrs at 30 °C with shaking (250 r.p.m.). 50 μL of the O/N culture was transferred into 450 μL fresh mineral medium and incubated for 72 hrs with the same conditions as described above. Cultures were centrifuged at 3,500 r.p.m., and supernatants were diluted 10-fold into mQ water before analysis on an Aminex HPX-87H ion exclusion column. Samples were analyzed for 45 min each at 60 °C and with a flow rate of 0.6 mL/min of 1 mM H_2_SO_4_. Quantifications of PCA and CCM were performed by comparison with the spectrum of standards ranging from 16 mg/L to 160 mg/L.

## Supporting information

Additional tables and figures can be found in the Supporting Information file.

## Acknowledgements

This work was supported by the Novo Nordisk Foundation. Authors would also like to thank Mette Christensen and Lars Schrübbers for technical advice related to HPLC.

## Author contributions

EDJ, MKJ, CCL, and JDK conceived the study. EDJ, FA, MBB and AAJ conducted all experimental work related to strain designs, constructions, and characterizations. EDJ, FA, and MBB performed all data analysis. EDJ and MKJ wrote the manuscript. All authors reviewed and approved the manuscript.

