## Supplementary Information for "Integrating continuous hypermutation with high-throughput screening for optimization of *cis,cis*-muconic acid production in yeast"

**Supplementary Table S1. p1 integration cassette sequence from pMB11.**

| Annotations | p1 homology  *KpAroY.B*  Kozak sequence  ORF10  *KpAroY.Ciso*  *URA3* |
| --- | --- |
| Sequence | ACTATAATATATGAATTACATTATTAATTTAAAACAACGGAATGCGTGCGATGAATTCCAAAGTCATGATTCAATTTCTTGAGCGAATTGTTCAGCAGTTCTCAAACCGTTCCATCTTCTAGCTTTGTGATAATCCAAACCGAATTGGTCCAAAACTCTGGTAACTATATGGTTGGTGATGTCATCAACGGTTTCTGGATGATTGTAATAAGCTGGCATTGGTGGAACCATAGCTACACCCATTCTAGACAAGGCCAACATGTTTTCCAAATGGATAGTAGACAATGGCATTTCTCTTGGGACCAAGACCAACTTTCTACCTTCTTTCAAAACAACATCAGCAGCTCTACCAACCAAACCTTCAGCATAACCAGCTCTAATACCAGCCAAGGTTTTCATAGAGCATGGAATAACGATCATACCATCAGTTCTGAAAGAACCAGAAGAAATAGTAGCAGCTTGATCAGCTGGAGAATGAGAAAAATCAGCCAAAGCAGCAACTTCTCTAGCAGTCCATGGAGTTTCCAATTCAATGGTGGTCTTAGCCCATTTAGACATAACCAAATGGGTTTCAACTTCTGGCATATCTCTCAAAGCTTGCAACAAAGCAACACCCAATGGAGCACCTGTAGCACCAGTCATACCGATGATCAACTTCATTGTTTTtttacatgtctatgagcttatcatatatttctacaAGCTCTCtgtagaaatatatgataagctcatagacatgtaaaAAAACAATGACCGCCCCAATCCAAGATTTGAGAGATGCTATTGCTTTGTTACAACAACACGACAATCAATACTTGGAAACCGATCATCCAGTTGATCCAAATGCTGAATTGGCTGGTGTTTACAGACATATTGGTGCTGGTGGTACTGTAAAAAGACCAACTAGAATTGGTCCAGCCATGATGTTCAACAACATTAAGGGTTATCCACACTCCAGAATCTTGGTTGGTATGCATGCTTCTAGACAAAGAGCAGCTTTGTTGTTGGGTTGTGAAGCTTCTCAATTGGCTTTGGAAGTTGGTAAAGCTGTTAAGAAACCAGTTGCTCCAGTTGTTGTTCCAGCTTCTTCTGCTCCATGTCAAGAACAAATTTTCTTGGCTGATGATCCAGACTTCGATTTGAGAACTTTGTTGCCAGCTCATACCAACACTCCAATTGATGCTGGTCCATTTTTTTGTTTGGGTTTGGCTTTAGCTTCTGATCCTGTTGATGCTTCTTTGACCGATGTTACCATTCATAGATTGTGCGTTCAAGGTAGAGATGAATTGTCTATGTTTTTGGCTGCCGGTAGACATATCGAAGTTTTTAGACAAAAAGCTGAAGCTGCTGGTAAGCCATTGCCAATTACTATTAACATGGGTTTAGATCCAGCCATCTACATTGGTGCTTGTTTTGAAGCTCCAACTACTCCATTTGGTTACAACGAATTGGGTGTTGCTGGTGCTTTGAGACAAAGACCAGTTGAATTGGTTCAAGGTGTTTCTGTTCCAGAAAAGGCTATTGCTAGAGCCGAAATAGTTATCGAAGGTGAATTATTGCCAGGTGTCAGAGTTAGAGAAGATCAACATACAAATTCCGGTCATGCTATGCCAGAATTTCCAGGTTATTGTGGTGGTGCTAATCCATCTTTGCCAGTTATTAAGGTTAAGGCCGTTACCATGAGAAACAACGCTATTTTACAAACTTTGGTCGGTCCAGGTGAAGAACATACAACTTTGGCTGGTTTGCCAACCGAAGCTTCTATTTGGAATGCTGTTGAAGCTGCAATTCCAGGTTTCTTGCAAAATGTTTATGCTCATACAGCTGGTGGTGGTAAGTTCTTGGGTATATTGCAAGTCAAGAAAAGACAACCAGCTGACGAAGGTAGACAAGGTCAAGCTGCTTTATTAGCTTTGGCTACTTACTCCGAATTGAAGAATATCATCTTGGTCGATGAAGATGTTGATATCTTCGATTCCGATGATATTTTGTGGGCTATGACTACTAGAATGCAAGGTGATGTTTCCATTACTACCATTCCAGGTATTAGAGGTCACCAATTAGATCCATCTCAAACCCCAGAATACTCCCCATCAATTAGAGGTAATGGTATCTCCTGTAAGACCATTTTCGATTGCACTGTTCCATGGGCTTTGAAGTCTCATTTTGAAAGAGCACCATTTGCTGACGTTGATCCTAGACCTTTTGCTCCAGAATATTTCGCTAGATTGGAAAAGAATCAAGGTTCCGCTAAGTCATGAATAATGAATTCCGCGTGCATTCTATATGAAAGTTTTTATAATAATTATAAAATGCATAAAGCTACATATAAGGAACGTGCTGCTACTCATCCTAGTCCTGTTGCTGCCAAGCTATTTAATATCATGCACGAAAAGCAAACAAACTTGTGTGCTTCATTGGATGTTCGTACCACCAAGGAATTACTGGAGTTAGTTGAAGCATTAGGTCCCAAAATTTGTTTACTAAAAACACATGTGGATATCTTGACTGATTTTTCCATGGAGGGCACAGTTAAGCCGCTAAAGGCATTATCCGCCAAGTACAATTTTTTACTCTTCGAAGACAGAAAATTTGCTGACATTGGTAATACAGTCAAATTGCAGTATTCTGCGGGTGTATACAGAATAGCAGAATGGGCAGACATTACGAATGCACACGGTGTGGTGGGCCCAGGTATTGTTAGCGGTTTGAAGCAGGCGGCAGAAGAAGTAACAAAGGAACCTAGAGGCCTTTTGATGTTAGCAGAATTGTCATGCAAGGGCTCCCTATCTACTGGAGAATATACTAAGGGTACTGTTGACATTGCGAAGAGCGACAAAGATTTTGTTATCGGCTTTATTGCTCAAAGAGACATGGGTGGAAGAGATGAAGGTTACGATTGGTTGATTATGACACCCGGTGTGGGTTTAGATGACAAGGGAGACGCATTGGGTCAACAGTATAGAACCGTGGATGATGTGGTCTCTACAGGATCTGACATTATTATTGTTGGAAGAGGACTATTTGCAAAGGGAAGGGATGCTAAGGTAGAGGGTGAACGTTACAGAAAAGCAGGCTGGGAAGCATATTTGAGAAGATGCGGCCAGCAAAACTAAGGTTATGCTTCTATTATAGAAAGAATAACTCTGGATTTAATGGAAATATATTCTATTAAAGGACTTAATGATATACCTAGAGATATAAAATTTAATATGGAAAAAATAAGACAAGAAAGATACAACCAAATGAAAGAAGCTCTAAATAGT |

**Supplementary Table S2. List of all sequences relevant to this study.**

| **Gene block** | **Mutation** | | **Sequence** |
| --- | --- | --- | --- |
| **General gene blocks** | | | |
| URA3-KO-90-mer | N/A | | ACCCAACTGCACAGAACAAAAACCTGCAGGAAACGAAGATAAATCAAAACTGTATTATAAGTAAATGCATGTATACTAAACTCACAAATT |
| gEC475 | N/A | | CTCGAGAGTACTATAATATATGAATTACATTATTAATTTAAAACAACGGAATGCGTGCGATCGCGTGCATTCTATATGAAAGTTTTTATAATAATTATAAAATGCATAAAGCTACATATAAGGAACGTGCTGCTACTCATCCTAGTCCTGTTGCTGCCAAGCTATTTAATATCATGCACGAAAAGCAAACAAACTTGTGTGCTTCATTGGATGTTCGTACCACCAAGGAATTACTGGAGTTAGTTGAAGCATTAGGTCCCAAAATTTGTTTACTAAAAACACATGTGGATATCTTGACTGATTTTTCCATGGAGGGCACAGTTAAGCCGCTAAAGGCATTATCCGCCAAGTACAATTTTTTACTCTTCGAAGACAGAAAATTTGCTGACATTGGTAATACAGTCAAATTGCAGTATTCTGCGGGTGTATACAGAATAGCAGAATGGGCAGACATTACGAATGCACACGGTGTGGTGGGCCCAGGTATTGTTAGCGGTTTGAAGCAGGCGGCAGAAGAAGTAACAAAGGAACCTAGAGGCCTTTTGATGTTAGCAGAATTGTCATGCAAGGGCTCCCTATCTACTGGAGAATATACTAAGGGTACTGTTGACATTGCGAAGAGCGACAAAGATTTTGTTATCGGCTTTATTGCTCAAAGAGACATGGGTGGAAGAGATGAAGGTTACGATTGGTTGATTATGACACCCGGTGTGGGTTTAGATGACAAGGGAGACGCATTGGGTCAACAGTATAGAACCGTGGATGATGTGGTCTCTACAGGATCTGACATTATTATTGTTGGAAGAGGACTATTTGCAAAGGGAAGGGATGCTAAGGTAGAGGGTGAACGTTACAGAAAAGCAGGCTGGGAAGCATATTTGAGAAGATGCGGCCAGCAAAACTAAGGTTATGCTTCTATTATAGAAAGAATAACTCTGGATTTAATGGAAATATATTCTATTAAAGGACTTAATGATATACCTAGAGATATAAAATTTAATATGGAAAAAATAAGACAAGAAAGATACAACCAAATGAAAGAAGCTCTAAATAGTACTTCTAGA |
| gEDJ12 | N/A | | ATCTGTCATAAAACAATGGAATTGAGACACTTGAGATACTTCGTTGCCGTTGTTGAAGAACAATCTTTTACAAAGGCTGCCGACAAGTTGTGTATTGCTCAACCACCATTATCCAGACAAATCCAAAACTTGGAAGAAGAATTGGGTATCCAATTATTGGAAAGAGGTTCCAGACCAGTTAAGACTACTCCAGAAGGTCATTTCTTTTACCAATACGCCATCAAGTTGCTAAGCAACGTTGATCAAATGGTCAGTATGACCAAGAGAATTGCCTCTGTCGAAAAGACCATTAGAATCGGTTTTGTTGGTTCCTTGTTGTTCGGTTTGTTGCCAAGAATTATCCACTTGTACAGACAAGCTCATCCAAACTTGAGAATCGAATTATACGAAATGGGTACTAAGGCTCAAACCGAAGCTTTGAAAGAAGGTAGAATTGACGCTGGTTTTGGTAGATTGAAGATTTCTGATCCAGCCATCAAGAGAACCTTGTTGAGAAACGAAAGATTGATGGTTGCTGTTCATGCTTCCCATCCATTGAATCAAATGAAGGATAAGGGTGTTCACTTGAACGATTTGATCGACGAAAAGATCTTGTTGTACCCATCTTCTCCAAAGCCAAACTTCTCTACTCATGTTATGAACATCTTCTCTGACCATGGTTTGGAACCTACCAAGATTAACGAAGTTAGAGAAGTCCAATTGGTCTTGGGTTTGGTTGCTGCTGGTGAAGGTATTTCATTGGTTCCAGCTTCTACCCAATCCATTCAATTATCCAACTTGTCCTACGTTCCATTATTAGATCCAGATGCTATTACCCCAATCTACATTGCTGTTAGAAACATGGAAGAATCCACCTACATCTACTCATTAAACGAAACCATCAGACAAATCCACGCCTACGAAGGTTTTACTGAACCACCGAATTGGTAAATCGCGTGCAT |
| **KpAroY.B and KpAroY.Ciso mutant gene blocks** | | | |
| KpAroY.Ciso_462T>C | Silent | | ATGACCGCCCCAATCCAAGATTTGAGAGATGCTATTGCTTTGTTACAACAACACGACAATCAATACTTGGAAACCGATCATCCAGTTGATCCAAATGCTGAATTGGCTGGTGTTTACAGACATATTGGTGCTGGTGGTACTGTAAAAAGACCAACTAGAATTGGTCCAGCCATGATGTTCAACAACATTAAGGGTTATCCACACTCCAGAATCTTGGTTGGTATGCATGCTTCTAGACAAAGAGCAGCTTTGTTGTTGGGTTGTGAAGCTTCTCAATTGGCTTTGGAAGTTGGTAAAGCTGTTAAGAAACCAGTTGCTCCAGTTGTTGTTCCAGCTTCTTCTGCTCCATGTCAAGAACAAATTTTCTTGGCTGATGATCCAGACTTCGATTTGAGAACTTTGTTGCCAGCTCATACCAACACTCCAATTGATGCTGGTCCATTTTTTTGTTTGGGTTTGGCCTTAGCTTCTGATCCTGTTGATGCTTCTTTGACCGATGTTACCATTCATAGATTGTGCGTTCAAGGTAGAGATGAATTGTCTATGTTTTTGGCTGCCGGTAGACATATCGAAGTTTTTAGACAAAAAGCTGAAGCTGCTGGTAAGCCATTGCCAATTACTATTAACATGGGTTTAGATCCAGCCATCTACATTGGTGCTTGTTTTGAAGCTCCAACTACTCCATTTGGTTACAACGAATTGGGTGTTGCTGGTGCTTTGAGACAAAGACCAGTTGAATTGGTTCAAGGTGTTTCTGTTCCAGAAAAGGCTATTGCTAGAGCCGAAATAGTTATCGAAGGTGAATTATTGCCAGGTGTCAGAGTTAGAGAAGATCAACATACAAATTCCGGTCATGCTATGCCAGAATTTCCAGGTTATTGTGGTGGTGCTAATCCATCTTTGCCAGTTATTAAGGTTAAGGCCGTTACCATGAGAAACAACGCTATTTTACAAACTTTGGTCGGTCCAGGTGAAGAACATACAACTTTGGCTGGTTTGCCAACCGAAGCTTCTATTTGGAATGCTGTTGAAGCTGCAATTCCAGGTTTCTTGCAAAATGTTTATGCTCATACAGCTGGTGGTGGTAAGTTCTTGGGTATATTGCAAGTCAAGAAAAGACAACCAGCTGACGAAGGTAGACAAGGTCAAGCTGCTTTATTAGCTTTGGCTACTTACTCCGAATTGAAGAATATCATCTTGGTCGATGAAGATGTTGATATCTTCGATTCCGATGATATTTTGTGGGCTATGACTACTAGAATGCAAGGTGATGTTTCCATTACTACCATTCCAGGTATTAGAGGTCACCAATTAGATCCATCTCAAACCCCAGAATACTCCCCATCAATTAGAGGTAATGGTATCTCCTGTAAGACCATTTTCGATTGCACTGTTCCATGGGCTTTGAAGTCTCATTTTGAAAGAGCACCATTTGCTGACGTTGATCCTAGACCTTTTGCTCCAGAATATTTCGCTAGATTGGAAAAGAATCAAGGTTCCGCTAAGTCATGA |
| KpAroY.Ciso_C150W | Missense | | ATGACCGCCCCAATCCAAGATTTGAGAGATGCTATTGCTTTGTTACAACAACACGACAATCAATACTTGGAAACCGATCATCCAGTTGATCCAAATGCTGAATTGGCTGGTGTTTACAGACATATTGGTGCTGGTGGTACTGTAAAAAGACCAACTAGAATTGGTCCAGCCATGATGTTCAACAACATTAAGGGTTATCCACACTCCAGAATCTTGGTTGGTATGCATGCTTCTAGACAAAGAGCAGCTTTGTTGTTGGGTTGTGAAGCTTCTCAATTGGCTTTGGAAGTTGGTAAAGCTGTTAAGAAACCAGTTGCTCCAGTTGTTGTTCCAGCTTCTTCTGCTCCATGTCAAGAACAAATTTTCTTGGCTGATGATCCAGACTTCGATTTGAGAACTTTGTTGCCAGCTCATACCAACACTCCAATTGATGCTGGTCCATTTTTTTGGTTGGGTTTGGCTTTAGCTTCTGATCCTGTTGATGCTTCTTTGACCGATGTTACCATTCATAGATTGTGCGTTCAAGGTAGAGATGAATTGTCTATGTTTTTGGCTGCCGGTAGACATATCGAAGTTTTTAGACAAAAAGCTGAAGCTGCTGGTAAGCCATTGCCAATTACTATTAACATGGGTTTAGATCCAGCCATCTACATTGGTGCTTGTTTTGAAGCTCCAACTACTCCATTTGGTTACAACGAATTGGGTGTTGCTGGTGCTTTGAGACAAAGACCAGTTGAATTGGTTCAAGGTGTTTCTGTTCCAGAAAAGGCTATTGCTAGAGCCGAAATAGTTATCGAAGGTGAATTATTGCCAGGTGTCAGAGTTAGAGAAGATCAACATACAAATTCCGGTCATGCTATGCCAGAATTTCCAGGTTATTGTGGTGGTGCTAATCCATCTTTGCCAGTTATTAAGGTTAAGGCCGTTACCATGAGAAACAACGCTATTTTACAAACTTTGGTCGGTCCAGGTGAAGAACATACAACTTTGGCTGGTTTGCCAACCGAAGCTTCTATTTGGAATGCTGTTGAAGCTGCAATTCCAGGTTTCTTGCAAAATGTTTATGCTCATACAGCTGGTGGTGGTAAGTTCTTGGGTATATTGCAAGTCAAGAAAAGACAACCAGCTGACGAAGGTAGACAAGGTCAAGCTGCTTTATTAGCTTTGGCTACTTACTCCGAATTGAAGAATATCATCTTGGTCGATGAAGATGTTGATATCTTCGATTCCGATGATATTTTGTGGGCTATGACTACTAGAATGCAAGGTGATGTTTCCATTACTACCATTCCAGGTATTAGAGGTCACCAATTAGATCCATCTCAAACCCCAGAATACTCCCCATCAATTAGAGGTAATGGTATCTCCTGTAAGACCATTTTCGATTGCACTGTTCCATGGGCTTTGAAGTCTCATTTTGAAAGAGCACCATTTGCTGACGTTGATCCTAGACCTTTTGCTCCAGAATATTTCGCTAGATTGGAAAAGAATCAAGGTTCCGCTAAGTCATGA |
| KpAroY.Ciso_780A>G | Silent | | ATGACCGCCCCAATCCAAGATTTGAGAGATGCTATTGCTTTGTTACAACAACACGACAATCAATACTTGGAAACCGATCATCCAGTTGATCCAAATGCTGAATTGGCTGGTGTTTACAGACATATTGGTGCTGGTGGTACTGTAAAAAGACCAACTAGAATTGGTCCAGCCATGATGTTCAACAACATTAAGGGTTATCCACACTCCAGAATCTTGGTTGGTATGCATGCTTCTAGACAAAGAGCAGCTTTGTTGTTGGGTTGTGAAGCTTCTCAATTGGCTTTGGAAGTTGGTAAAGCTGTTAAGAAACCAGTTGCTCCAGTTGTTGTTCCAGCTTCTTCTGCTCCATGTCAAGAACAAATTTTCTTGGCTGATGATCCAGACTTCGATTTGAGAACTTTGTTGCCAGCTCATACCAACACTCCAATTGATGCTGGTCCATTTTTTTGTTTGGGTTTGGCTTTAGCTTCTGATCCTGTTGATGCTTCTTTGACCGATGTTACCATTCATAGATTGTGCGTTCAAGGTAGAGATGAATTGTCTATGTTTTTGGCTGCCGGTAGACATATCGAAGTTTTTAGACAAAAAGCTGAAGCTGCTGGTAAGCCATTGCCAATTACTATTAACATGGGTTTAGATCCAGCCATCTACATTGGTGCTTGTTTTGAAGCTCCAACTACTCCATTTGGTTACAACGAATTGGGTGTTGCTGGTGCTTTGAGACAAAGACCAGTTGAATTGGTTCAAGGTGTTTCTGTTCCAGAAAAGGCTATTGCTAGGGCCGAAATAGTTATCGAAGGTGAATTATTGCCAGGTGTCAGAGTTAGAGAAGATCAACATACAAATTCCGGTCATGCTATGCCAGAATTTCCAGGTTATTGTGGTGGTGCTAATCCATCTTTGCCAGTTATTAAGGTTAAGGCCGTTACCATGAGAAACAACGCTATTTTACAAACTTTGGTCGGTCCAGGTGAAGAACATACAACTTTGGCTGGTTTGCCAACCGAAGCTTCTATTTGGAATGCTGTTGAAGCTGCAATTCCAGGTTTCTTGCAAAATGTTTATGCTCATACAGCTGGTGGTGGTAAGTTCTTGGGTATATTGCAAGTCAAGAAAAGACAACCAGCTGACGAAGGTAGACAAGGTCAAGCTGCTTTATTAGCTTTGGCTACTTACTCCGAATTGAAGAATATCATCTTGGTCGATGAAGATGTTGATATCTTCGATTCCGATGATATTTTGTGGGCTATGACTACTAGAATGCAAGGTGATGTTTCCATTACTACCATTCCAGGTATTAGAGGTCACCAATTAGATCCATCTCAAACCCCAGAATACTCCCCATCAATTAGAGGTAATGGTATCTCCTGTAAGACCATTTTCGATTGCACTGTTCCATGGGCTTTGAAGTCTCATTTTGAAAGAGCACCATTTGCTGACGTTGATCCTAGACCTTTTGCTCCAGAATATTTCGCTAGATTGGAAAAGAATCAAGGTTCCGCTAAGTCATGA |
| KpAroY.Ciso_700T>C | Silent | | ATGACCGCCCCAATCCAAGATTTGAGAGATGCTATTGCTTTGTTACAACAACACGACAATCAATACTTGGAAACCGATCATCCAGTTGATCCAAATGCTGAATTGGCTGGTGTTTACAGACATATTGGTGCTGGTGGTACTGTAAAAAGACCAACTAGAATTGGTCCAGCCATGATGTTCAACAACATTAAGGGTTATCCACACTCCAGAATCTTGGTTGGTATGCATGCTTCTAGACAAAGAGCAGCTTTGTTGTTGGGTTGTGAAGCTTCTCAATTGGCTTTGGAAGTTGGTAAAGCTGTTAAGAAACCAGTTGCTCCAGTTGTTGTTCCAGCTTCTTCTGCTCCATGTCAAGAACAAATTTTCTTGGCTGATGATCCAGACTTCGATTTGAGAACTTTGTTGCCAGCTCATACCAACACTCCAATTGATGCTGGTCCATTTTTTTGTTTGGGTTTGGCTTTAGCTTCTGATCCTGTTGATGCTTCTTTGACCGATGTTACCATTCATAGATTGTGCGTTCAAGGTAGAGATGAATTGTCTATGTTTTTGGCTGCCGGTAGACATATCGAAGTTTTTAGACAAAAAGCTGAAGCTGCTGGTAAGCCATTGCCAATTACTATTAACATGGGTTTAGATCCAGCCATCTACATTGGTGCTTGTTTTGAAGCTCCAACTACTCCATTTGGTTACAACGAACTGGGTGTTGCTGGTGCTTTGAGACAAAGACCAGTTGAATTGGTTCAAGGTGTTTCTGTTCCAGAAAAGGCTATTGCTAGAGCCGAAATAGTTATCGAAGGTGAATTATTGCCAGGTGTCAGAGTTAGAGAAGATCAACATACAAATTCCGGTCATGCTATGCCAGAATTTCCAGGTTATTGTGGTGGTGCTAATCCATCTTTGCCAGTTATTAAGGTTAAGGCCGTTACCATGAGAAACAACGCTATTTTACAAACTTTGGTCGGTCCAGGTGAAGAACATACAACTTTGGCTGGTTTGCCAACCGAAGCTTCTATTTGGAATGCTGTTGAAGCTGCAATTCCAGGTTTCTTGCAAAATGTTTATGCTCATACAGCTGGTGGTGGTAAGTTCTTGGGTATATTGCAAGTCAAGAAAAGACAACCAGCTGACGAAGGTAGACAAGGTCAAGCTGCTTTATTAGCTTTGGCTACTTACTCCGAATTGAAGAATATCATCTTGGTCGATGAAGATGTTGATATCTTCGATTCCGATGATATTTTGTGGGCTATGACTACTAGAATGCAAGGTGATGTTTCCATTACTACCATTCCAGGTATTAGAGGTCACCAATTAGATCCATCTCAAACCCCAGAATACTCCCCATCAATTAGAGGTAATGGTATCTCCTGTAAGACCATTTTCGATTGCACTGTTCCATGGGCTTTGAAGTCTCATTTTGAAAGAGCACCATTTGCTGACGTTGATCCTAGACCTTTTGCTCCAGAATATTTCGCTAGATTGGAAAAGAATCAAGGTTCCGCTAAGTCATGA |
| KpAroY.Ciso_L134F | Missense | | ATGACCGCCCCAATCCAAGATTTGAGAGATGCTATTGCTTTGTTACAACAACACGACAATCAATACTTGGAAACCGATCATCCAGTTGATCCAAATGCTGAATTGGCTGGTGTTTACAGACATATTGGTGCTGGTGGTACTGTAAAAAGACCAACTAGAATTGGTCCAGCCATGATGTTCAACAACATTAAGGGTTATCCACACTCCAGAATCTTGGTTGGTATGCATGCTTCTAGACAAAGAGCAGCTTTGTTGTTGGGTTGTGAAGCTTCTCAATTGGCTTTGGAAGTTGGTAAAGCTGTTAAGAAACCAGTTGCTCCAGTTGTTGTTCCAGCTTCTTCTGCTCCATGTCAAGAACAAATTTTCTTGGCTGATGATCCAGACTTCGATTTGAGAACTTTTTTGCCAGCTCATACCAACACTCCAATTGATGCTGGTCCATTTTTTTGTTTGGGTTTGGCTTTAGCTTCTGATCCTGTTGATGCTTCTTTGACCGATGTTACCATTCATAGATTGTGCGTTCAAGGTAGAGATGAATTGTCTATGTTTTTGGCTGCCGGTAGACATATCGAAGTTTTTAGACAAAAAGCTGAAGCTGCTGGTAAGCCATTGCCAATTACTATTAACATGGGTTTAGATCCAGCCATCTACATTGGTGCTTGTTTTGAAGCTCCAACTACTCCATTTGGTTACAACGAATTGGGTGTTGCTGGTGCTTTGAGACAAAGACCAGTTGAATTGGTTCAAGGTGTTTCTGTTCCAGAAAAGGCTATTGCTAGAGCCGAAATAGTTATCGAAGGTGAATTATTGCCAGGTGTCAGAGTTAGAGAAGATCAACATACAAATTCCGGTCATGCTATGCCAGAATTTCCAGGTTATTGTGGTGGTGCTAATCCATCTTTGCCAGTTATTAAGGTTAAGGCCGTTACCATGAGAAACAACGCTATTTTACAAACTTTGGTCGGTCCAGGTGAAGAACATACAACTTTGGCTGGTTTGCCAACCGAAGCTTCTATTTGGAATGCTGTTGAAGCTGCAATTCCAGGTTTCTTGCAAAATGTTTATGCTCATACAGCTGGTGGTGGTAAGTTCTTGGGTATATTGCAAGTCAAGAAAAGACAACCAGCTGACGAAGGTAGACAAGGTCAAGCTGCTTTATTAGCTTTGGCTACTTACTCCGAATTGAAGAATATCATCTTGGTCGATGAAGATGTTGATATCTTCGATTCCGATGATATTTTGTGGGCTATGACTACTAGAATGCAAGGTGATGTTTCCATTACTACCATTCCAGGTATTAGAGGTCACCAATTAGATCCATCTCAAACCCCAGAATACTCCCCATCAATTAGAGGTAATGGTATCTCCTGTAAGACCATTTTCGATTGCACTGTTCCATGGGCTTTGAAGTCTCATTTTGAAAGAGCACCATTTGCTGACGTTGATCCTAGACCTTTTGCTCCAGAATATTTCGCTAGATTGGAAAAGAATCAAGGTTCCGCTAAGTCATGA |
| KpAroY.Ciso_A308T | Missense | | ATGACCGCCCCAATCCAAGATTTGAGAGATGCTATTGCTTTGTTACAACAACACGACAATCAATACTTGGAAACCGATCATCCAGTTGATCCAAATGCTGAATTGGCTGGTGTTTACAGACATATTGGTGCTGGTGGTACTGTAAAAAGACCAACTAGAATTGGTCCAGCCATGATGTTCAACAACATTAAGGGTTATCCACACTCCAGAATCTTGGTTGGTATGCATGCTTCTAGACAAAGAGCAGCTTTGTTGTTGGGTTGTGAAGCTTCTCAATTGGCTTTGGAAGTTGGTAAAGCTGTTAAGAAACCAGTTGCTCCAGTTGTTGTTCCAGCTTCTTCTGCTCCATGTCAAGAACAAATTTTCTTGGCTGATGATCCAGACTTCGATTTGAGAACTTTGTTGCCAGCTCATACCAACACTCCAATTGATGCTGGTCCATTTTTTTGTTTGGGTTTGGCTTTAGCTTCTGATCCTGTTGATGCTTCTTTGACCGATGTTACCATTCATAGATTGTGCGTTCAAGGTAGAGATGAATTGTCTATGTTTTTGGCTGCCGGTAGACATATCGAAGTTTTTAGACAAAAAGCTGAAGCTGCTGGTAAGCCATTGCCAATTACTATTAACATGGGTTTAGATCCAGCCATCTACATTGGTGCTTGTTTTGAAGCTCCAACTACTCCATTTGGTTACAACGAATTGGGTGTTGCTGGTGCTTTGAGACAAAGACCAGTTGAATTGGTTCAAGGTGTTTCTGTTCCAGAAAAGGCTATTGCTAGAGCCGAAATAGTTATCGAAGGTGAATTATTGCCAGGTGTCAGAGTTAGAGAAGATCAACATACAAATTCCGGTCATGCTATGCCAGAATTTCCAGGTTATTGTGGTGGTGCTAATCCATCTTTGCCAGTTATTAAGGTTAAGACCGTTACCATGAGAAACAACGCTATTTTACAAACTTTGGTCGGTCCAGGTGAAGAACATACAACTTTGGCTGGTTTGCCAACCGAAGCTTCTATTTGGAATGCTGTTGAAGCTGCAATTCCAGGTTTCTTGCAAAATGTTTATGCTCATACAGCTGGTGGTGGTAAGTTCTTGGGTATATTGCAAGTCAAGAAAAGACAACCAGCTGACGAAGGTAGACAAGGTCAAGCTGCTTTATTAGCTTTGGCTACTTACTCCGAATTGAAGAATATCATCTTGGTCGATGAAGATGTTGATATCTTCGATTCCGATGATATTTTGTGGGCTATGACTACTAGAATGCAAGGTGATGTTTCCATTACTACCATTCCAGGTATTAGAGGTCACCAATTAGATCCATCTCAAACCCCAGAATACTCCCCATCAATTAGAGGTAATGGTATCTCCTGTAAGACCATTTTCGATTGCACTGTTCCATGGGCTTTGAAGTCTCATTTTGAAAGAGCACCATTTGCTGACGTTGATCCTAGACCTTTTGCTCCAGAATATTTCGCTAGATTGGAAAAGAATCAAGGTTCCGCTAAGTCATGA |
| KpAroY.Ciso_E474V | Missense | | ATGACCGCCCCAATCCAAGATTTGAGAGATGCTATTGCTTTGTTACAACAACACGACAATCAATACTTGGAAACCGATCATCCAGTTGATCCAAATGCTGAATTGGCTGGTGTTTACAGACATATTGGTGCTGGTGGTACTGTAAAAAGACCAACTAGAATTGGTCCAGCCATGATGTTCAACAACATTAAGGGTTATCCACACTCCAGAATCTTGGTTGGTATGCATGCTTCTAGACAAAGAGCAGCTTTGTTGTTGGGTTGTGAAGCTTCTCAATTGGCTTTGGAAGTTGGTAAAGCTGTTAAGAAACCAGTTGCTCCAGTTGTTGTTCCAGCTTCTTCTGCTCCATGTCAAGAACAAATTTTCTTGGCTGATGATCCAGACTTCGATTTGAGAACTTTGTTGCCAGCTCATACCAACACTCCAATTGATGCTGGTCCATTTTTTTGTTTGGGTTTGGCTTTAGCTTCTGATCCTGTTGATGCTTCTTTGACCGATGTTACCATTCATAGATTGTGCGTTCAAGGTAGAGATGAATTGTCTATGTTTTTGGCTGCCGGTAGACATATCGAAGTTTTTAGACAAAAAGCTGAAGCTGCTGGTAAGCCATTGCCAATTACTATTAACATGGGTTTAGATCCAGCCATCTACATTGGTGCTTGTTTTGAAGCTCCAACTACTCCATTTGGTTACAACGAATTGGGTGTTGCTGGTGCTTTGAGACAAAGACCAGTTGAATTGGTTCAAGGTGTTTCTGTTCCAGAAAAGGCTATTGCTAGAGCCGAAATAGTTATCGAAGGTGAATTATTGCCAGGTGTCAGAGTTAGAGAAGATCAACATACAAATTCCGGTCATGCTATGCCAGAATTTCCAGGTTATTGTGGTGGTGCTAATCCATCTTTGCCAGTTATTAAGGTTAAGGCCGTTACCATGAGAAACAACGCTATTTTACAAACTTTGGTCGGTCCAGGTGAAGAACATACAACTTTGGCTGGTTTGCCAACCGAAGCTTCTATTTGGAATGCTGTTGAAGCTGCAATTCCAGGTTTCTTGCAAAATGTTTATGCTCATACAGCTGGTGGTGGTAAGTTCTTGGGTATATTGCAAGTCAAGAAAAGACAACCAGCTGACGAAGGTAGACAAGGTCAAGCTGCTTTATTAGCTTTGGCTACTTACTCCGAATTGAAGAATATCATCTTGGTCGATGAAGATGTTGATATCTTCGATTCCGATGATATTTTGTGGGCTATGACTACTAGAATGCAAGGTGATGTTTCCATTACTACCATTCCAGGTATTAGAGGTCACCAATTAGATCCATCTCAAACCCCAGAATACTCCCCATCAATTAGAGGTAATGGTATCTCCTGTAAGACCATTTTCGATTGCACTGTTCCATGGGCTTTGAAGTCTCATTTTGTAAGAGCACCATTTGCTGACGTTGATCCTAGACCTTTTGCTCCAGAATATTTCGCTAGATTGGAAAAGAATCAAGGTTCCGCTAAGTCATGA |
| KpAroY.Ciso_522T>C | Silent | | ATGACCGCCCCAATCCAAGATTTGAGAGATGCTATTGCTTTGTTACAACAACACGACAATCAATACTTGGAAACCGATCATCCAGTTGATCCAAATGCTGAATTGGCTGGTGTTTACAGACATATTGGTGCTGGTGGTACTGTAAAAAGACCAACTAGAATTGGTCCAGCCATGATGTTCAACAACATTAAGGGTTATCCACACTCCAGAATCTTGGTTGGTATGCATGCTTCTAGACAAAGAGCAGCTTTGTTGTTGGGTTGTGAAGCTTCTCAATTGGCTTTGGAAGTTGGTAAAGCTGTTAAGAAACCAGTTGCTCCAGTTGTTGTTCCAGCTTCTTCTGCTCCATGTCAAGAACAAATTTTCTTGGCTGATGATCCAGACTTCGATTTGAGAACTTTGTTGCCAGCTCATACCAACACTCCAATTGATGCTGGTCCATTTTTTTGTTTGGGTTTGGCTTTAGCTTCTGATCCTGTTGATGCTTCTTTGACCGATGTTACCATTCATAGATTGTGCGTCCAAGGTAGAGATGAATTGTCTATGTTTTTGGCTGCCGGTAGACATATCGAAGTTTTTAGACAAAAAGCTGAAGCTGCTGGTAAGCCATTGCCAATTACTATTAACATGGGTTTAGATCCAGCCATCTACATTGGTGCTTGTTTTGAAGCTCCAACTACTCCATTTGGTTACAACGAATTGGGTGTTGCTGGTGCTTTGAGACAAAGACCAGTTGAATTGGTTCAAGGTGTTTCTGTTCCAGAAAAGGCTATTGCTAGAGCCGAAATAGTTATCGAAGGTGAATTATTGCCAGGTGTCAGAGTTAGAGAAGATCAACATACAAATTCCGGTCATGCTATGCCAGAATTTCCAGGTTATTGTGGTGGTGCTAATCCATCTTTGCCAGTTATTAAGGTTAAGGCCGTTACCATGAGAAACAACGCTATTTTACAAACTTTGGTCGGTCCAGGTGAAGAACATACAACTTTGGCTGGTTTGCCAACCGAAGCTTCTATTTGGAATGCTGTTGAAGCTGCAATTCCAGGTTTCTTGCAAAATGTTTATGCTCATACAGCTGGTGGTGGTAAGTTCTTGGGTATATTGCAAGTCAAGAAAAGACAACCAGCTGACGAAGGTAGACAAGGTCAAGCTGCTTTATTAGCTTTGGCTACTTACTCCGAATTGAAGAATATCATCTTGGTCGATGAAGATGTTGATATCTTCGATTCCGATGATATTTTGTGGGCTATGACTACTAGAATGCAAGGTGATGTTTCCATTACTACCATTCCAGGTATTAGAGGTCACCAATTAGATCCATCTCAAACCCCAGAATACTCCCCATCAATTAGAGGTAATGGTATCTCCTGTAAGACCATTTTCGATTGCACTGTTCCATGGGCTTTGAAGTCTCATTTTGAAAGAGCACCATTTGCTGACGTTGATCCTAGACCTTTTGCTCCAGAATATTTCGCTAGATTGGAAAAGAATCAAGGTTCCGCTAAGTCATGA |
| KpAroY.Ciso_V401A | Missense | | ATGACCGCCCCAATCCAAGATTTGAGAGATGCTATTGCTTTGTTACAACAACACGACAATCAATACTTGGAAACCGATCATCCAGTTGATCCAAATGCTGAATTGGCTGGTGTTTACAGACATATTGGTGCTGGTGGTACTGTAAAAAGACCAACTAGAATTGGTCCAGCCATGATGTTCAACAACATTAAGGGTTATCCACACTCCAGAATCTTGGTTGGTATGCATGCTTCTAGACAAAGAGCAGCTTTGTTGTTGGGTTGTGAAGCTTCTCAATTGGCTTTGGAAGTTGGTAAAGCTGTTAAGAAACCAGTTGCTCCAGTTGTTGTTCCAGCTTCTTCTGCTCCATGTCAAGAACAAATTTTCTTGGCTGATGATCCAGACTTCGATTTGAGAACTTTGTTGCCAGCTCATACCAACACTCCAATTGATGCTGGTCCATTTTTTTGTTTGGGTTTGGCTTTAGCTTCTGATCCTGTTGATGCTTCTTTGACCGATGTTACCATTCATAGATTGTGCGTTCAAGGTAGAGATGAATTGTCTATGTTTTTGGCTGCCGGTAGACATATCGAAGTTTTTAGACAAAAAGCTGAAGCTGCTGGTAAGCCATTGCCAATTACTATTAACATGGGTTTAGATCCAGCCATCTACATTGGTGCTTGTTTTGAAGCTCCAACTACTCCATTTGGTTACAACGAATTGGGTGTTGCTGGTGCTTTGAGACAAAGACCAGTTGAATTGGTTCAAGGTGTTTCTGTTCCAGAAAAGGCTATTGCTAGAGCCGAAATAGTTATCGAAGGTGAATTATTGCCAGGTGTCAGAGTTAGAGAAGATCAACATACAAATTCCGGTCATGCTATGCCAGAATTTCCAGGTTATTGTGGTGGTGCTAATCCATCTTTGCCAGTTATTAAGGTTAAGGCCGTTACCATGAGAAACAACGCTATTTTACAAACTTTGGTCGGTCCAGGTGAAGAACATACAACTTTGGCTGGTTTGCCAACCGAAGCTTCTATTTGGAATGCTGTTGAAGCTGCAATTCCAGGTTTCTTGCAAAATGTTTATGCTCATACAGCTGGTGGTGGTAAGTTCTTGGGTATATTGCAAGTCAAGAAAAGACAACCAGCTGACGAAGGTAGACAAGGTCAAGCTGCTTTATTAGCTTTGGCTACTTACTCCGAATTGAAGAATATCATCTTGGCCGATGAAGATGTTGATATCTTCGATTCCGATGATATTTTGTGGGCTATGACTACTAGAATGCAAGGTGATGTTTCCATTACTACCATTCCAGGTATTAGAGGTCACCAATTAGATCCATCTCAAACCCCAGAATACTCCCCATCAATTAGAGGTAATGGTATCTCCTGTAAGACCATTTTCGATTGCACTGTTCCATGGGCTTTGAAGTCTCATTTTGAAAGAGCACCATTTGCTGACGTTGATCCTAGACCTTTTGCTCCAGAATATTTCGCTAGATTGGAAAAGAATCAAGGTTCCGCTAAGTCATGA |
| KpAroY.Ciso_966T>C | Silent | | ATGACCGCCCCAATCCAAGATTTGAGAGATGCTATTGCTTTGTTACAACAACACGACAATCAATACTTGGAAACCGATCATCCAGTTGATCCAAATGCTGAATTGGCTGGTGTTTACAGACATATTGGTGCTGGTGGTACTGTAAAAAGACCAACTAGAATTGGTCCAGCCATGATGTTCAACAACATTAAGGGTTATCCACACTCCAGAATCTTGGTTGGTATGCATGCTTCTAGACAAAGAGCAGCTTTGTTGTTGGGTTGTGAAGCTTCTCAATTGGCTTTGGAAGTTGGTAAAGCTGTTAAGAAACCAGTTGCTCCAGTTGTTGTTCCAGCTTCTTCTGCTCCATGTCAAGAACAAATTTTCTTGGCTGATGATCCAGACTTCGATTTGAGAACTTTGTTGCCAGCTCATACCAACACTCCAATTGATGCTGGTCCATTTTTTTGTTTGGGTTTGGCTTTAGCTTCTGATCCTGTTGATGCTTCTTTGACCGATGTTACCATTCATAGATTGTGCGTTCAAGGTAGAGATGAATTGTCTATGTTTTTGGCTGCCGGTAGACATATCGAAGTTTTTAGACAAAAAGCTGAAGCTGCTGGTAAGCCATTGCCAATTACTATTAACATGGGTTTAGATCCAGCCATCTACATTGGTGCTTGTTTTGAAGCTCCAACTACTCCATTTGGTTACAACGAATTGGGTGTTGCTGGTGCTTTGAGACAAAGACCAGTTGAATTGGTTCAAGGTGTTTCTGTTCCAGAAAAGGCTATTGCTAGAGCCGAAATAGTTATCGAAGGTGAATTATTGCCAGGTGTCAGAGTTAGAGAAGATCAACATACAAATTCCGGTCATGCTATGCCAGAATTTCCAGGTTATTGTGGTGGTGCTAATCCATCTTTGCCAGTTATTAAGGTTAAGGCCGTTACCATGAGAAACAACGCTATTTTACAAACTTTGGTCGGCCCAGGTGAAGAACATACAACTTTGGCTGGTTTGCCAACCGAAGCTTCTATTTGGAATGCTGTTGAAGCTGCAATTCCAGGTTTCTTGCAAAATGTTTATGCTCATACAGCTGGTGGTGGTAAGTTCTTGGGTATATTGCAAGTCAAGAAAAGACAACCAGCTGACGAAGGTAGACAAGGTCAAGCTGCTTTATTAGCTTTGGCTACTTACTCCGAATTGAAGAATATCATCTTGGTCGATGAAGATGTTGATATCTTCGATTCCGATGATATTTTGTGGGCTATGACTACTAGAATGCAAGGTGATGTTTCCATTACTACCATTCCAGGTATTAGAGGTCACCAATTAGATCCATCTCAAACCCCAGAATACTCCCCATCAATTAGAGGTAATGGTATCTCCTGTAAGACCATTTTCGATTGCACTGTTCCATGGGCTTTGAAGTCTCATTTTGAAAGAGCACCATTTGCTGACGTTGATCCTAGACCTTTTGCTCCAGAATATTTCGCTAGATTGGAAAAGAATCAAGGTTCCGCTAAGTCATGA |
| KpAroY.Ciso_F222L | Missense | | ATGACCGCCCCAATCCAAGATTTGAGAGATGCTATTGCTTTGTTACAACAACACGACAATCAATACTTGGAAACCGATCATCCAGTTGATCCAAATGCTGAATTGGCTGGTGTTTACAGACATATTGGTGCTGGTGGTACTGTAAAAAGACCAACTAGAATTGGTCCAGCCATGATGTTCAACAACATTAAGGGTTATCCACACTCCAGAATCTTGGTTGGTATGCATGCTTCTAGACAAAGAGCAGCTTTGTTGTTGGGTTGTGAAGCTTCTCAATTGGCTTTGGAAGTTGGTAAAGCTGTTAAGAAACCAGTTGCTCCAGTTGTTGTTCCAGCTTCTTCTGCTCCATGTCAAGAACAAATTTTCTTGGCTGATGATCCAGACTTCGATTTGAGAACTTTGTTGCCAGCTCATACCAACACTCCAATTGATGCTGGTCCATTTTTTTGTTTGGGTTTGGCTTTAGCTTCTGATCCTGTTGATGCTTCTTTGACCGATGTTACCATTCATAGATTGTGCGTTCAAGGTAGAGATGAATTGTCTATGTTTTTGGCTGCCGGTAGACATATCGAAGTTTTTAGACAAAAAGCTGAAGCTGCTGGTAAGCCATTGCCAATTACTATTAACATGGGTTTAGATCCAGCCATCTACATTGGTGCTTGTTTGGAAGCTCCAACTACTCCATTTGGTTACAACGAATTGGGTGTTGCTGGTGCTTTGAGACAAAGACCAGTTGAATTGGTTCAAGGTGTTTCTGTTCCAGAAAAGGCTATTGCTAGAGCCGAAATAGTTATCGAAGGTGAATTATTGCCAGGTGTCAGAGTTAGAGAAGATCAACATACAAATTCCGGTCATGCTATGCCAGAATTTCCAGGTTATTGTGGTGGTGCTAATCCATCTTTGCCAGTTATTAAGGTTAAGGCCGTTACCATGAGAAACAACGCTATTTTACAAACTTTGGTCGGTCCAGGTGAAGAACATACAACTTTGGCTGGTTTGCCAACCGAAGCTTCTATTTGGAATGCTGTTGAAGCTGCAATTCCAGGTTTCTTGCAAAATGTTTATGCTCATACAGCTGGTGGTGGTAAGTTCTTGGGTATATTGCAAGTCAAGAAAAGACAACCAGCTGACGAAGGTAGACAAGGTCAAGCTGCTTTATTAGCTTTGGCTACTTACTCCGAATTGAAGAATATCATCTTGGTCGATGAAGATGTTGATATCTTCGATTCCGATGATATTTTGTGGGCTATGACTACTAGAATGCAAGGTGATGTTTCCATTACTACCATTCCAGGTATTAGAGGTCACCAATTAGATCCATCTCAAACCCCAGAATACTCCCCATCAATTAGAGGTAATGGTATCTCCTGTAAGACCATTTTCGATTGCACTGTTCCATGGGCTTTGAAGTCTCATTTTGAAAGAGCACCATTTGCTGACGTTGATCCTAGACCTTTTGCTCCAGAATATTTCGCTAGATTGGAAAAGAATCAAGGTTCCGCTAAGTCATGA |
| KpAroY.Ciso_408A>G | Silent | | ATGACCGCCCCAATCCAAGATTTGAGAGATGCTATTGCTTTGTTACAACAACACGACAATCAATACTTGGAAACCGATCATCCAGTTGATCCAAATGCTGAATTGGCTGGTGTTTACAGACATATTGGTGCTGGTGGTACTGTAAAAAGACCAACTAGAATTGGTCCAGCCATGATGTTCAACAACATTAAGGGTTATCCACACTCCAGAATCTTGGTTGGTATGCATGCTTCTAGACAAAGAGCAGCTTTGTTGTTGGGTTGTGAAGCTTCTCAATTGGCTTTGGAAGTTGGTAAAGCTGTTAAGAAACCAGTTGCTCCAGTTGTTGTTCCAGCTTCTTCTGCTCCATGTCAAGAACAAATTTTCTTGGCTGATGATCCAGACTTCGATTTGAGAACTTTGTTGCCGGCTCATACCAACACTCCAATTGATGCTGGTCCATTTTTTTGTTTGGGTTTGGCTTTAGCTTCTGATCCTGTTGATGCTTCTTTGACCGATGTTACCATTCATAGATTGTGCGTTCAAGGTAGAGATGAATTGTCTATGTTTTTGGCTGCCGGTAGACATATCGAAGTTTTTAGACAAAAAGCTGAAGCTGCTGGTAAGCCATTGCCAATTACTATTAACATGGGTTTAGATCCAGCCATCTACATTGGTGCTTGTTTTGAAGCTCCAACTACTCCATTTGGTTACAACGAATTGGGTGTTGCTGGTGCTTTGAGACAAAGACCAGTTGAATTGGTTCAAGGTGTTTCTGTTCCAGAAAAGGCTATTGCTAGAGCCGAAATAGTTATCGAAGGTGAATTATTGCCAGGTGTCAGAGTTAGAGAAGATCAACATACAAATTCCGGTCATGCTATGCCAGAATTTCCAGGTTATTGTGGTGGTGCTAATCCATCTTTGCCAGTTATTAAGGTTAAGGCCGTTACCATGAGAAACAACGCTATTTTACAAACTTTGGTCGGTCCAGGTGAAGAACATACAACTTTGGCTGGTTTGCCAACCGAAGCTTCTATTTGGAATGCTGTTGAAGCTGCAATTCCAGGTTTCTTGCAAAATGTTTATGCTCATACAGCTGGTGGTGGTAAGTTCTTGGGTATATTGCAAGTCAAGAAAAGACAACCAGCTGACGAAGGTAGACAAGGTCAAGCTGCTTTATTAGCTTTGGCTACTTACTCCGAATTGAAGAATATCATCTTGGTCGATGAAGATGTTGATATCTTCGATTCCGATGATATTTTGTGGGCTATGACTACTAGAATGCAAGGTGATGTTTCCATTACTACCATTCCAGGTATTAGAGGTCACCAATTAGATCCATCTCAAACCCCAGAATACTCCCCATCAATTAGAGGTAATGGTATCTCCTGTAAGACCATTTTCGATTGCACTGTTCCATGGGCTTTGAAGTCTCATTTTGAAAGAGCACCATTTGCTGACGTTGATCCTAGACCTTTTGCTCCAGAATATTTCGCTAGATTGGAAAAGAATCAAGGTTCCGCTAAGTCATGA |
| KpAroY.Ciso_1047T>C | Silent | | ATGACCGCCCCAATCCAAGATTTGAGAGATGCTATTGCTTTGTTACAACAACACGACAATCAATACTTGGAAACCGATCATCCAGTTGATCCAAATGCTGAATTGGCTGGTGTTTACAGACATATTGGTGCTGGTGGTACTGTAAAAAGACCAACTAGAATTGGTCCAGCCATGATGTTCAACAACATTAAGGGTTATCCACACTCCAGAATCTTGGTTGGTATGCATGCTTCTAGACAAAGAGCAGCTTTGTTGTTGGGTTGTGAAGCTTCTCAATTGGCTTTGGAAGTTGGTAAAGCTGTTAAGAAACCAGTTGCTCCAGTTGTTGTTCCAGCTTCTTCTGCTCCATGTCAAGAACAAATTTTCTTGGCTGATGATCCAGACTTCGATTTGAGAACTTTGTTGCCAGCTCATACCAACACTCCAATTGATGCTGGTCCATTTTTTTGTTTGGGTTTGGCTTTAGCTTCTGATCCTGTTGATGCTTCTTTGACCGATGTTACCATTCATAGATTGTGCGTTCAAGGTAGAGATGAATTGTCTATGTTTTTGGCTGCCGGTAGACATATCGAAGTTTTTAGACAAAAAGCTGAAGCTGCTGGTAAGCCATTGCCAATTACTATTAACATGGGTTTAGATCCAGCCATCTACATTGGTGCTTGTTTTGAAGCTCCAACTACTCCATTTGGTTACAACGAATTGGGTGTTGCTGGTGCTTTGAGACAAAGACCAGTTGAATTGGTTCAAGGTGTTTCTGTTCCAGAAAAGGCTATTGCTAGAGCCGAAATAGTTATCGAAGGTGAATTATTGCCAGGTGTCAGAGTTAGAGAAGATCAACATACAAATTCCGGTCATGCTATGCCAGAATTTCCAGGTTATTGTGGTGGTGCTAATCCATCTTTGCCAGTTATTAAGGTTAAGGCCGTTACCATGAGAAACAACGCTATTTTACAAACTTTGGTCGGTCCAGGTGAAGAACATACAACTTTGGCTGGTTTGCCAACCGAAGCTTCTATTTGGAATGCTGTTGAAGCTGCAATTCCAGGCTTCTTGCAAAATGTTTATGCTCATACAGCTGGTGGTGGTAAGTTCTTGGGTATATTGCAAGTCAAGAAAAGACAACCAGCTGACGAAGGTAGACAAGGTCAAGCTGCTTTATTAGCTTTGGCTACTTACTCCGAATTGAAGAATATCATCTTGGTCGATGAAGATGTTGATATCTTCGATTCCGATGATATTTTGTGGGCTATGACTACTAGAATGCAAGGTGATGTTTCCATTACTACCATTCCAGGTATTAGAGGTCACCAATTAGATCCATCTCAAACCCCAGAATACTCCCCATCAATTAGAGGTAATGGTATCTCCTGTAAGACCATTTTCGATTGCACTGTTCCATGGGCTTTGAAGTCTCATTTTGAAAGAGCACCATTTGCTGACGTTGATCCTAGACCTTTTGCTCCAGAATATTTCGCTAGATTGGAAAAGAATCAAGGTTCCGCTAAGTCATGA |
| KpAroY.Ciso_P444L | Missense | | ATGACCGCCCCAATCCAAGATTTGAGAGATGCTATTGCTTTGTTACAACAACACGACAATCAATACTTGGAAACCGATCATCCAGTTGATCCAAATGCTGAATTGGCTGGTGTTTACAGACATATTGGTGCTGGTGGTACTGTAAAAAGACCAACTAGAATTGGTCCAGCCATGATGTTCAACAACATTAAGGGTTATCCACACTCCAGAATCTTGGTTGGTATGCATGCTTCTAGACAAAGAGCAGCTTTGTTGTTGGGTTGTGAAGCTTCTCAATTGGCTTTGGAAGTTGGTAAAGCTGTTAAGAAACCAGTTGCTCCAGTTGTTGTTCCAGCTTCTTCTGCTCCATGTCAAGAACAAATTTTCTTGGCTGATGATCCAGACTTCGATTTGAGAACTTTGTTGCCAGCTCATACCAACACTCCAATTGATGCTGGTCCATTTTTTTGTTTGGGTTTGGCTTTAGCTTCTGATCCTGTTGATGCTTCTTTGACCGATGTTACCATTCATAGATTGTGCGTTCAAGGTAGAGATGAATTGTCTATGTTTTTGGCTGCCGGTAGACATATCGAAGTTTTTAGACAAAAAGCTGAAGCTGCTGGTAAGCCATTGCCAATTACTATTAACATGGGTTTAGATCCAGCCATCTACATTGGTGCTTGTTTTGAAGCTCCAACTACTCCATTTGGTTACAACGAATTGGGTGTTGCTGGTGCTTTGAGACAAAGACCAGTTGAATTGGTTCAAGGTGTTTCTGTTCCAGAAAAGGCTATTGCTAGAGCCGAAATAGTTATCGAAGGTGAATTATTGCCAGGTGTCAGAGTTAGAGAAGATCAACATACAAATTCCGGTCATGCTATGCCAGAATTTCCAGGTTATTGTGGTGGTGCTAATCCATCTTTGCCAGTTATTAAGGTTAAGGCCGTTACCATGAGAAACAACGCTATTTTACAAACTTTGGTCGGTCCAGGTGAAGAACATACAACTTTGGCTGGTTTGCCAACCGAAGCTTCTATTTGGAATGCTGTTGAAGCTGCAATTCCAGGTTTCTTGCAAAATGTTTATGCTCATACAGCTGGTGGTGGTAAGTTCTTGGGTATATTGCAAGTCAAGAAAAGACAACCAGCTGACGAAGGTAGACAAGGTCAAGCTGCTTTATTAGCTTTGGCTACTTACTCCGAATTGAAGAATATCATCTTGGTCGATGAAGATGTTGATATCTTCGATTCCGATGATATTTTGTGGGCTATGACTACTAGAATGCAAGGTGATGTTTCCATTACTACCATTCCAGGTATTAGAGGTCACCAATTAGATCCATCTCAAACCCTAGAATACTCCCCATCAATTAGAGGTAATGGTATCTCCTGTAAGACCATTTTCGATTGCACTGTTCCATGGGCTTTGAAGTCTCATTTTGAAAGAGCACCATTTGCTGACGTTGATCCTAGACCTTTTGCTCCAGAATATTTCGCTAGATTGGAAAAGAATCAAGGTTCCGCTAAGTCATGA |
| KpAroY.Ciso_N497S | Missense | | ATGACCGCCCCAATCCAAGATTTGAGAGATGCTATTGCTTTGTTACAACAACACGACAATCAATACTTGGAAACCGATCATCCAGTTGATCCAAATGCTGAATTGGCTGGTGTTTACAGACATATTGGTGCTGGTGGTACTGTAAAAAGACCAACTAGAATTGGTCCAGCCATGATGTTCAACAACATTAAGGGTTATCCACACTCCAGAATCTTGGTTGGTATGCATGCTTCTAGACAAAGAGCAGCTTTGTTGTTGGGTTGTGAAGCTTCTCAATTGGCTTTGGAAGTTGGTAAAGCTGTTAAGAAACCAGTTGCTCCAGTTGTTGTTCCAGCTTCTTCTGCTCCATGTCAAGAACAAATTTTCTTGGCTGATGATCCAGACTTCGATTTGAGAACTTTGTTGCCAGCTCATACCAACACTCCAATTGATGCTGGTCCATTTTTTTGTTTGGGTTTGGCTTTAGCTTCTGATCCTGTTGATGCTTCTTTGACCGATGTTACCATTCATAGATTGTGCGTTCAAGGTAGAGATGAATTGTCTATGTTTTTGGCTGCCGGTAGACATATCGAAGTTTTTAGACAAAAAGCTGAAGCTGCTGGTAAGCCATTGCCAATTACTATTAACATGGGTTTAGATCCAGCCATCTACATTGGTGCTTGTTTTGAAGCTCCAACTACTCCATTTGGTTACAACGAATTGGGTGTTGCTGGTGCTTTGAGACAAAGACCAGTTGAATTGGTTCAAGGTGTTTCTGTTCCAGAAAAGGCTATTGCTAGAGCCGAAATAGTTATCGAAGGTGAATTATTGCCAGGTGTCAGAGTTAGAGAAGATCAACATACAAATTCCGGTCATGCTATGCCAGAATTTCCAGGTTATTGTGGTGGTGCTAATCCATCTTTGCCAGTTATTAAGGTTAAGGCCGTTACCATGAGAAACAACGCTATTTTACAAACTTTGGTCGGTCCAGGTGAAGAACATACAACTTTGGCTGGTTTGCCAACCGAAGCTTCTATTTGGAATGCTGTTGAAGCTGCAATTCCAGGTTTCTTGCAAAATGTTTATGCTCATACAGCTGGTGGTGGTAAGTTCTTGGGTATATTGCAAGTCAAGAAAAGACAACCAGCTGACGAAGGTAGACAAGGTCAAGCTGCTTTATTAGCTTTGGCTACTTACTCCGAATTGAAGAATATCATCTTGGTCGATGAAGATGTTGATATCTTCGATTCCGATGATATTTTGTGGGCTATGACTACTAGAATGCAAGGTGATGTTTCCATTACTACCATTCCAGGTATTAGAGGTCACCAATTAGATCCATCTCAAACCCCAGAATACTCCCCATCAATTAGAGGTAATGGTATCTCCTGTAAGACCATTTTCGATTGCACTGTTCCATGGGCTTTGAAGTCTCATTTTGAAAGAGCACCATTTGCTGACGTTGATCCTAGACCTTTTGCTCCAGAATATTTCGCTAGATTGGAAAAGAGTCAAGGTTCCGCTAAGTCATGA |
| KpAroY.Ciso_666T>del | deletion | | ATGACCGCCCCAATCCAAGATTTGAGAGATGCTATTGCTTTGTTACAACAACACGACAATCAATACTTGGAAACCGATCATCCAGTTGATCCAAATGCTGAATTGGCTGGTGTTTACAGACATATTGGTGCTGGTGGTACTGTAAAAAGACCAACTAGAATTGGTCCAGCCATGATGTTCAACAACATTAAGGGTTATCCACACTCCAGAATCTTGGTTGGTATGCATGCTTCTAGACAAAGAGCAGCTTTGTTGTTGGGTTGTGAAGCTTCTCAATTGGCTTTGGAAGTTGGTAAAGCTGTTAAGAAACCAGTTGCTCCAGTTGTTGTTCCAGCTTCTTCTGCTCCATGTCAAGAACAAATTTTCTTGGCTGATGATCCAGACTTCGATTTGAGAACTTTGTTGCCAGCTCATACCAACACTCCAATTGATGCTGGTCCATTTTTTTGTTTGGGTTTGGCTTTAGCTTCTGATCCTGTTGATGCTTCTTTGACCGATGTTACCATTCATAGATTGTGCGTTCAAGGTAGAGATGAATTGTCTATGTTTTTGGCTGCCGGTAGACATATCGAAGTTTTTAGACAAAAAGCTGAAGCTGCTGGTAAGCCATTGCCAATTACTATTAACATGGGTTTAGATCCAGCCATCTACATTGGTGCTTGTTTGAAGCTCCAACTACTCCATTTGGTTACAACGAATTGGGTGTTGCTGGTGCTTTGAGACAAAGACCAGTTGAATTGGTTCAAGGTGTTTCTGTTCCAGAAAAGGCTATTGCTAGAGCCGAAATAGTTATCGAAGGTGAATTATTGCCAGGTGTCAGAGTTAGAGAAGATCAACATACAAATTCCGGTCATGCTATGCCAGAATTTCCAGGTTATTGTGGTGGTGCTAATCCATCTTTGCCAGTTATTAAGGTTAAGGCCGTTACCATGAGAAACAACGCTATTTTACAAACTTTGGTCGGTCCAGGTGAAGAACATACAACTTTGGCTGGTTTGCCAACCGAAGCTTCTATTTGGAATGCTGTTGAAGCTGCAATTCCAGGTTTCTTGCAAAATGTTTATGCTCATACAGCTGGTGGTGGTAAGTTCTTGGGTATATTGCAAGTCAAGAAAAGACAACCAGCTGACGAAGGTAGACAAGGTCAAGCTGCTTTATTAGCTTTGGCTACTTACTCCGAATTGAAGAATATCATCTTGGTCGATGAAGATGTTGATATCTTCGATTCCGATGATATTTTGTGGGCTATGACTACTAGAATGCAAGGTGATGTTTCCATTACTACCATTCCAGGTATTAGAGGTCACCAATTAGATCCATCTCAAACCCCAGAATACTCCCCATCAATTAGAGGTAATGGTATCTCCTGTAAGACCATTTTCGATTGCACTGTTCCATGGGCTTTGAAGTCTCATTTTGAAAGAGCACCATTTGCTGACGTTGATCCTAGACCTTTTGCTCCAGAATATTTCGCTAGATTGGAAAAGAATCAAGGTTCCGCTAAGTCATGA |
| KpAroY.B_T157A | Missense | | ATGAAGTTGATCATCGGTATGACTGGTGCTACAGGTGCTCCATTGGGTGTTGCTTTGTTGCAAGCTTTGAGAGATATGCCAGAAGTTGAAACCCATTTGGTTATGTCTAAATGGGCTAAGACCACCATTGAATTGGAAACTCCATGGACTGCTAGAGAAGTTGCTGCTTTGGCTGATTTTTCTCATTCTCCAGCTGATCAAGCTGCTACTATTTCTTCTGGTTCTTTCAGAACTGATGGTATGATCGTTATTCCATGCTCTATGAAAACCTTGGCTGGTATTAGAGCTGGTTATGCTGAAGGTTTGGTTGGTAGAGCTGCTGATGTTGTTTTGAAAGAAGGTAGAAAGTTGGTCTTGGTCCCAAGAGAAATGCCATTGTCTACTATCCATTTGGAAAACATGTTGGCCTTGTCTAGAATGGGTGTAGCTATGGTTCCACCAATGCCAGCTTATTACAATCATCCAGAAGCCGTTGATGACATCACCAACCATATAGTTACCAGAGTTTTGGACCAATTCGGTTTGGATTATCACAAAGCTAGAAGATGGAACGGTTTGAGAACTGCTGAACAATTCGCTCAAGAAATTGAATCATGA |
| KpAroY.B_P146T | Missense | | ATGAAGTTGATCATCGGTATGACTGGTGCTACAGGTGCTCCATTGGGTGTTGCTTTGTTGCAAGCTTTGAGAGATATGCCAGAAGTTGAAACCCATTTGGTTATGTCTAAATGGGCTAAGACCACCATTGAATTGGAAACTCCATGGACTGCTAGAGAAGTTGCTGCTTTGGCTGATTTTTCTCATTCTCCAGCTGATCAAGCTGCTACTATTTCTTCTGGTTCTTTCAGAACTGATGGTATGATCGTTATTCCATGCTCTATGAAAACCTTGGCTGGTATTAGAGCTGGTTATGCTGAAGGTTTGGTTGGTAGAGCTGCTGATGTTGTTTTGAAAGAAGGTAGAAAGTTGGTCTTGGTCCCAAGAGAAATGCCATTGTCTACTATCCATTTGGAAAACATGTTGGCCTTGTCTAGAATGGGTGTAGCTATGGTTACACCAATGCCAGCTTATTACAATCATCCAGAAACCGTTGATGACATCACCAACCATATAGTTACCAGAGTTTTGGACCAATTCGGTTTGGATTATCACAAAGCTAGAAGATGGAACGGTTTGAGAACTGCTGAACAATTCGCTCAAGAAATTGAATCATGA |
| KpAroY.B_315A>G | Silent | | ATGAAGTTGATCATCGGTATGACTGGTGCTACAGGTGCTCCATTGGGTGTTGCTTTGTTGCAAGCTTTGAGAGATATGCCAGAAGTTGAAACCCATTTGGTTATGTCTAAATGGGCTAAGACCACCATTGAATTGGAAACTCCATGGACTGCTAGAGAAGTTGCTGCTTTGGCTGATTTTTCTCATTCTCCAGCTGATCAAGCTGCTACTATTTCTTCTGGTTCTTTCAGAACTGATGGTATGATCGTTATTCCATGCTCTATGAAAACCTTGGCTGGTATTAGAGCTGGTTATGCTGAAGGTTTGGTTGGTAGGGCTGCTGATGTTGTTTTGAAAGAAGGTAGAAAGTTGGTCTTGGTCCCAAGAGAAATGCCATTGTCTACTATCCATTTGGAAAACATGTTGGCCTTGTCTAGAATGGGTGTAGCTATGGTTCCACCAATGCCAGCTTATTACAATCATCCAGAAACCGTTGATGACATCACCAACCATATAGTTACCAGAGTTTTGGACCAATTCGGTTTGGATTATCACAAAGCTAGAAGATGGAACGGTTTGAGAACTGCTGAACAATTCGCTCAAGAAATTGAATCATGA |
| KpAroY.B_33A>G | Silent | | ATGAAGTTGATCATCGGTATGACTGGTGCTACGGGTGCTCCATTGGGTGTTGCTTTGTTGCAAGCTTTGAGAGATATGCCAGAAGTTGAAACCCATTTGGTTATGTCTAAATGGGCTAAGACCACCATTGAATTGGAAACTCCATGGACTGCTAGAGAAGTTGCTGCTTTGGCTGATTTTTCTCATTCTCCAGCTGATCAAGCTGCTACTATTTCTTCTGGTTCTTTCAGAACTGATGGTATGATCGTTATTCCATGCTCTATGAAAACCTTGGCTGGTATTAGAGCTGGTTATGCTGAAGGTTTGGTTGGTAGAGCTGCTGATGTTGTTTTGAAAGAAGGTAGAAAGTTGGTCTTGGTCCCAAGAGAAATGCCATTGTCTACTATCCATTTGGAAAACATGTTGGCCTTGTCTAGAATGGGTGTAGCTATGGTTCCACCAATGCCAGCTTATTACAATCATCCAGAAACCGTTGATGACATCACCAACCATATAGTTACCAGAGTTTTGGACCAATTCGGTTTGGATTATCACAAAGCTAGAAGATGGAACGGTTTGAGAACTGCTGAACAATTCGCTCAAGAAATTGAATCATGA |
| KpAroY.B_555T>del | Deletion | ATGAAGTTGATCATCGGTATGACTGGTGCTACAGGTGCTCCATTGGGTGTTGCTTTGTTGCAAGCTTTGAGAGATATGCCAGAAGTTGAAACCCATTTGGTTATGTCTAAATGGGCTAAGACCACCATTGAATTGGAAACTCCATGGACTGCTAGAGAAGTTGCTGCTTTGGCTGATTTTTCTCATTCTCCAGCTGATCAAGCTGCTACTATTTCTTCTGGTTCTTTCAGAACTGATGGTATGATCGTTATTCCATGCTCTATGAAAACCTTGGCTGGTATTAGAGCTGGTTATGCTGAAGGTTTGGTTGGTAGAGCTGCTGATGTTGTTTTGAAAGAAGGTAGAAAGTTGGTCTTGGTCCCAAGAGAAATGCCATTGTCTACTATCCATTTGGAAAACATGTTGGCCTTGTCTAGAATGGGTGTAGCTATGGTTCCACCAATGCCAGCTTATTACAATCATCCAGAAACCGTTGATGACATCACCAACCATATAGTTACCAGAGTTTTGGACCAATTCGGTTTGGATTATCACAAAGCTAGAAGATGGAACGGTTGAGAACTGCTGAACAATTCGCTCAAGAAATTGAATCATGA | |
| KpAroY.B_N184Y | Missense | ATGAAGTTGATCATCGGTATGACTGGTGCTACAGGTGCTCCATTGGGTGTTGCTTTGTTGCAAGCTTTGAGAGATATGCCAGAAGTTGAAACCCATTTGGTTATGTCTAAATGGGCTAAGACCACCATTGAATTGGAAACTCCATGGACTGCTAGAGAAGTTGCTGCTTTGGCTGATTTTTCTCATTCTCCAGCTGATCAAGCTGCTACTATTTCTTCTGGTTCTTTCAGAACTGATGGTATGATCGTTATTCCATGCTCTATGAAAACCTTGGCTGGTATTAGAGCTGGTTATGCTGAAGGTTTGGTTGGTAGAGCTGCTGATGTTGTTTTGAAAGAAGGTAGAAAGTTGGTCTTGGTCCCAAGAGAAATGCCATTGTCTACTATCCATTTGGAAAACATGTTGGCCTTGTCTAGAATGGGTGTAGCTATGGTTCCACCAATGCCAGCTTATTACAATCATCCAGAAACCGTTGATGACATCACCAACCATATAGTTACCAGAGTTTTGGACCAATTCGGTTTGGATTATCACAAAGCTAGAAGATGGTACGGTTTGAGAACTGCTGAACAATTCGCTCAAGAAATTGAATCATGA | |

**Supplementary Table S3. List of oligonucleotides used in this study.**

| **Oligo** | **5' modification** | **Sequence** | **Description** |
| --- | --- | --- | --- |
| ORP1 | N/A | CAATCTGGCGGCTTGAGTTC | F_XII-5-up |
| ORP2 | N/A | TGAGAACTGCTGAACAATTCGCTCAAGAAATTGAATCATGAGTAGATACGTTGTTGACAC | R_XII-5-up |
| ORP3 | N/A | AGAAAGCATAGCAATCTAATCTAAGTTTTAATTACAAAAAACAATGAAGTTGATCATCGG | F_KpAroY.B_Int |
| ORP4 | N/A | AAGAAATTCGCTTATTTAGAAGTGTCAACAACGTATCTACTCATGATTCAATTTCTTGAG | R_KpAroY.B_Int |
| ORP5 | N/A | GTAGCACCAGTCATACCGATGATCAACTTCATTGTTTTTTGTAATTAAAACTTAGATTAG | F_TEF1pro |
| ORP6 | N/A | agctccagcttttgttcccttcgagtcatgtaattagttaGCACACACCATAGCTTC | R_TEF1pro |
| ORP7 | N/A | aactaattacatgactcgaagggaacaaaagctggagctATAAAAAACACGCTTTTTCAG | F_TDH3pro |
| ORP8 | N/A | TCTCTCAAATCTTGGATTGGGGCGGTCATTGTTTTTTTGTTTGTTTATGTGTGTTTATTC | R_TDH3pro |
| ORP9 | N/A | ACTTAGTTTCGAATAAACACACATAAACAAACAAAAAAACAATGACCGCCCCAATCCAAG | F_KpAroY.Ciso_Int |
| ORP10 | N/A | ACTTCAGGTTGTCTAACTCCTTCCTTTTCGGTTAGAGCGGATTCATGACTTAGCGGAACC | R_KpAroY.Ciso_Int |
| ORP11 | N/A | CGCTAGATTGGAAAAGAATCAAGGTTCCGCTAAGTCATGAATCCGCTCTAACCGAAAAGG | F_XII-5-down |
| ORP12 | N/A | CTGCGATACCTTTTGTGATGG | R_XII-5-down |
| EDJ382 | N/A | TTAGCAGAATTGTCATGCAAGTTTTAGAGCTAGAAATAGCAAG | F_pCfB3050 |
| EDJ383 | 5/PHOS/ | GATCATTTATCTTTCACTGCGGAGAAG | R_pCfB3050 |
| EDJ386 | N/A | CGTACGGAATTCAGAGTACTGACAATAAAAAGATTCTTG | F_p1237 |
| EDJ387 | N/A | CTAGTCCTGCAGGGGTAACGCCAGG | R_p1237 |
| EDJ388 | N/A | CGTACGCTGCAGAACGACATTACTATATATATAATATAGGAAG | F_TRP1 |
| EDJ389 | N/A | CGTACGGAATTCAGGCAAGTGCACAAACAATAC | R_TRP1 |
| EDJ390 | N/A | ATAATCTCGAGAGTACTATAATATATGAATTAC | F_gEC475 |
| EDJ391 | N/A | ATAATTCTAGAAGTACTATTTAGAGCTTCTTTC | R_gEC475 |
| EDJ413 | N/A | CGTGCGAUGAATTCCAAAGTCATGATTCAATTTCTTGAGCGAATTG | F_KpAroY.B_p1 |
| EDJ414 | N/A | AGAGAGCUgtagaaatatatgataagctcatagacatgtaaaAAAACAATGAAGTTGATC | R_KpAroY.B_p1 |
| EDJ415 | N/A | AGCTCTCUgtagaaatatatgataagctcatagacatgtaaaAAAACAATGACCGCCCCA | F_KpAroY.Ciso_p1 |
| EDJ416 | N/A | CACGCGAUGAATTCATTATTCATGACTTAGCGGAACCTTGATTC | R_KpAroY.Ciso_p1 |
| EDJ432 | N/A | ATGACAGAUCGCTGGATATGCCTAGAAATGC | F_pCfB2764 |
| EDJ433 | N/A | ATCGCGTGCAUTCATCCGCTCTAACC | R_pCfB2764 |
| EDJ434 | N/A | ATCTGTCAUAAAACAATGGAATTGAGACACTTGAG | F_gEDJ12 |
| EDJ435 | N/A | ATGCACGCGAUTTACCAATTCGGTGGTTCAGTAAAACC | R_gEDJ12 |
| p1F | N/A | GCGATGAATTCCAAAGTC | Amplicon sequencing of p1 |
| p1R | N/A | CCTTATATGTAGCTTTATGC | Amplicon sequencing of p1 |
| p1S | N/A | CAATCCAAGATTTGAGAGATGC | Amplicon sequencing of p1 |

**Supplementary Table S4. List of plasmids used in this study.**

| **Plasmid** | **Content** | **Genetic marker** | **Antibiotic marker** | **Origin of replication in yeast** | **Backbone vector** | **Reference** |
| --- | --- | --- | --- | --- | --- | --- |
| pRS414-TEF1p-Cas9-CYC1t | TEF1p-Cas9-CYC1t | *TRP1* | *AmpR* | CEN6/ARS4 | N/A | (DiCarlo *et al.*, 2013) |
| p1237 | pX-4-loxP-SpHIS5-ScTkl1<-PTDH3-PTEF1->KpAroY.D | *SpHIS5* | *AmpR* | N/A | N/A | (Skjoedt *et al.*, 2016) |
| p1239 | pXI-1-LoxP-KlLEU2-PaAroZ<-PTDH3-PTEF1->CaCatA | *KILEU2* | *AmpR* | N/A | N/A | (Skjoedt *et al.*, 2016) |
| p1241 | pTY4-URA3-KpAroY.B<-PTDH3-PTEF1->KpAroY.Ciso | *KIURA3* | *AmpR* | N/A | N/A | (Skjoedt *et al.*, 2016) |
| pCfB2553 | pXII-4-LoxP-HphMXsyn-209bp_CYC1p_BenO_T1->yEGFP | *HphMXsyn* | *AmpR* | N/A | N/A | (Skjoedt *et al.*, 2016) |
| pEDJ515 | pX-3-LoxP-KanMXsyn-REV1p->BenM_MP17_D08 | *KanMXsyn* | *AmpR* | N/A | pCfB2764 | This study |
| pCfB2909 | XII-5-MarkerFree | No | *AmpR* | N/A | N/A | (Jessop-Fabre *et al.*, 2016) |
| pCfB3050 | SNR52p-gRNA_XII-5-SUP4t | *Nourseothricin(clonNat)* | *AmpR* | 2μ | N/A | (Jessop-Fabre *et al.*, 2016) |
| pEDJ366 | SNR52p-gRNA_URA3-SUP4t | *Nourseothricin(clonNat)* | *AmpR* | 2μ | pCfB3050 | This study |
| pEDJ371 | pX-4-loxP-TRP1-ScTkl1<-PTDH3-PTEF1->KpAroY.D | *TRP1* | *AmpR* | N/A | p1237 | This study |
| ZZ-Ec475 | p1 recombination cassette that integrates *Tm*HisA and URA3 in place of ORFs 1-4 on the orthogonal plasmid (pGKL1) | *URA3* | *AmpR* | N/A | N/A | (Zhong *et al.*, 2020) |
| pMB10 | ORF_URA3; USER cloning site; p1 homology arms | *URA3* | *AmpR* | N/A | ZZ-Ec475 | This study |
| pMB11 | URA3-KpAroY.B<-ORF10-ORF10->KpAroY.Ciso | *URA3* | *AmpR* | N/A | pMB10 | This study |
| AR-Ec633 | HIS3-TPDNAP1 (L477V, L640Y, I777K, W814N) | *HIS3* | *KanR* | CEN6/ARS4 | N/A | (Ravikumar *et al.*, 2018) |

**Supplementary Table S5. List of strains used in this study.**

| **Strain** | **Genotype** | **Plasmids** | **Parental strain** | **Reference** |
| --- | --- | --- | --- | --- |
| CEN.PK2-1C | MATa; his3D1; leu2-3_112; ura3-52; trp1-289; MAL2-8c; SUC2 | N/A | N/A | EUROSCARF |
| Sc-62 | MATa; his3D1; leu2-3_112; Δura3; trp1-289; MAL2-8c; SUC2 | pRS414-TEF1p-Cas9-CYC1t | CEN.PK2-1C | This study |
| Sc-67 | MATa; his3D1; leu2-3_112; Δura3; trp1-289; MAL2-8c; SUC2; XI-1-LoxP-KlLEU2-PaAroZ<-PTDH3-PTEF1->CaCatA | N/A | Sc-62 | This study |
| Sc-68 | MATa; his3D1; leu2-3_112; Δura3; trp1-289; MAL2-8c; SUC2; XI-1-LoxP-KlLEU2-PaAroZ<-PTDH3-PTEF1->CaCatA; X-4-loxP-TRP1-ScTkl1<-PTDH3-PTEF1->KpAroY.D | N/A | Sc-67 | This study |
| Sc-78 | MATa; his3D1; leu2-3_112; Δura3; trp1-289; MAL2-8c; SUC2; XI-1-LoxP-KlLEU2-PaAroZ<-PTDH3-PTEF1->CaCatA; X-4-loxP-TRP1-ScTkl1<-PTDH3-PTEF1->KpAroY.D; pXII-4-LoxP-HphMXsyn-209bp_CYC1p_BenO_T1->yEGFP; pX-3-KanMXsyn-REV1p::BenM-MP17_D08 | N/A | Sc-68 | This study |
| Sc-79 | MATa; his3D1; leu2-3_112; Δura3; trp1-289; MAL2-8c; SUC2; XI-1-LoxP-KlLEU2-PaAroZ<-PTDH3-PTEF1->CaCatA; X-4-loxP-TRP1-ScTkl1<-PTDH3-PTEF1->KpAroY.D; pXII-4-LoxP-HphMXsyn-209bp_CYC1p_BenO_T1->yEGFP; pX-3-KanMXsyn-REV1p::BenM-MP17_D08 | AR-Ec633 | Sc-78 | This study |
| F102-2 | MATa can1 his4-519 leu2-3, 112 ρ0 | p1 + p2 from *K. lactis* | N/A | ATCC (Catalog #200585) |
| Sc-93 | MATa can1 his4-519 leu2-3, 112 ρ0 | p1(pMB11 insert) and p2 | F102-2 | This study |
| Sc-105 | MATa; his3D1; leu2-3_112; Δura3; trp1-289; MAL2-8c; SUC2; XI-1-LoxP-KlLEU2-PaAroZ<-PTDH3-PTEF1->CaCatA; X-4-loxP-TRP1-ScTkl1<-PTDH3-PTEF1->KpAroY.D; pXII-4-LoxP-HphMXsyn-209bp_CYC1p_BenO_T1->yEGFP; pX-3-KanMXsyn-REV1p::BenM-MP17_D08 | AR-Ec633; p1(pMB11 insert) and p2 | Sc-79 and Sc-93 | This study |
| Sc-194 | MATa; his3D1; leu2-3_112; Δura3; trp1-289; MAL2-8c; SUC2; XI-1-LoxP-KlLEU2-PaAroZ<-PTDH3-PTEF1->CaCatA; X-4-loxP-TRP1-ScTkl1<-PTDH3-PTEF1->KpAroY.D; pXII-4-LoxP-HphMXsyn-209bp_CYC1p_BenO_T1->yEGFP; pX-3-KanMXsyn-REV1p::BenM-MP17_D08; XII-5-KpAroY.B<-PTEF1-PTDH3->KpAroY.Ciso | N/A | Sc-78 | This study |
| KpAroY.B_P146T | MATa; his3D1; leu2-3_112; Δura3; trp1-289; MAL2-8c; SUC2; XI-1-LoxP-KlLEU2-PaAroZ<-PTDH3-PTEF1->CaCatA; X-4-loxP-TRP1-ScTkl1<-PTDH3-PTEF1->KpAroY.D; pXII-4-LoxP-HphMXsyn-209bp_CYC1p_BenO_T1->yEGFP; pX-3-KanMXsyn-REV1p::BenM-MP17_D08; XII-5-KpAroY.B_P146T<-PTEF1-PTDH3->KpAroY.Ciso | N/A | Sc-78 | This study |
| KpAroB.Ciso_T157A | MATa; his3D1; leu2-3_112; Δura3; trp1-289; MAL2-8c; SUC2; XI-1-LoxP-KlLEU2-PaAroZ<-PTDH3-PTEF1->CaCatA; X-4-loxP-TRP1-ScTkl1<-PTDH3-PTEF1->KpAroY.D; pXII-4-LoxP-HphMXsyn-209bp_CYC1p_BenO_T1->yEGFP; pX-3-KanMXsyn-REV1p::BenM-MP17_D08; XII-5-KpAroY.B<-PTEF1-PTDH3->KpAroY.Ciso_T157A | N/A | Sc-78 | This study |
| KpAroY.Ciso_E474V | MATa; his3D1; leu2-3_112; Δura3; trp1-289; MAL2-8c; SUC2; XI-1-LoxP-KlLEU2-PaAroZ<-PTDH3-PTEF1->CaCatA; X-4-loxP-TRP1-ScTkl1<-PTDH3-PTEF1->KpAroY.D; pXII-4-LoxP-HphMXsyn-209bp_CYC1p_BenO_T1->yEGFP; pX-3-KanMXsyn-REV1p::BenM-MP17_D08; XII-5-KpAroY.B<-PTEF1-PTDH3->KpAroY.Ciso_E474V | N/A | Sc-78 | This study |
| KpAroY.Ciso_V401A | MATa; his3D1; leu2-3_112; Δura3; trp1-289; MAL2-8c; SUC2; XI-1-LoxP-KlLEU2-PaAroZ<-PTDH3-PTEF1->CaCatA; X-4-loxP-TRP1-ScTkl1<-PTDH3-PTEF1->KpAroY.D; pXII-4-LoxP-HphMXsyn-209bp_CYC1p_BenO_T1->yEGFP; pX-3-KanMXsyn-REV1p::BenM-MP17_D08; XII-5-KpAroY.B<-PTEF1-PTDH3->KpAroY.Ciso_V401A | N/A | Sc-78 | This study |
| KpAroY.Ciso_F222L | MATa; his3D1; leu2-3_112; Δura3; trp1-289; MAL2-8c; SUC2; XI-1-LoxP-KlLEU2-PaAroZ<-PTDH3-PTEF1->CaCatA; X-4-loxP-TRP1-ScTkl1<-PTDH3-PTEF1->KpAroY.D; pXII-4-LoxP-HphMXsyn-209bp_CYC1p_BenO_T1->yEGFP; pX-3-KanMXsyn-REV1p::BenM-MP17_D08; XII-5-KpAroY.B<-PTEF1-PTDH3->KpAroY.Ciso_F222L | N/A | Sc-78 | This study |
| KpAroY.Ciso_N497S | MATa; his3D1; leu2-3_112; Δura3; trp1-289; MAL2-8c; SUC2; XI-1-LoxP-KlLEU2-PaAroZ<-PTDH3-PTEF1->CaCatA; X-4-loxP-TRP1-ScTkl1<-PTDH3-PTEF1->KpAroY.D; pXII-4-LoxP-HphMXsyn-209bp_CYC1p_BenO_T1->yEGFP; pX-3-KanMXsyn-REV1p::BenM-MP17_D08; XII-5-KpAroY.B<-PTEF1-PTDH3->KpAroY.Ciso_N497S | N/A | Sc-78 | This study |
| KpAroY.Ciso_A308T | MATa; his3D1; leu2-3_112; Δura3; trp1-289; MAL2-8c; SUC2; XI-1-LoxP-KlLEU2-PaAroZ<-PTDH3-PTEF1->CaCatA; X-4-loxP-TRP1-ScTkl1<-PTDH3-PTEF1->KpAroY.D; pXII-4-LoxP-HphMXsyn-209bp_CYC1p_BenO_T1->yEGFP; pX-3-KanMXsyn-REV1p::BenM-MP17_D08; XII-5-KpAroY.B<-PTEF1-PTDH3->KpAroY.Ciso_A308T | N/A | Sc-78 | This study |
| KpAroY.Ciso_C150W | MATa; his3D1; leu2-3_112; Δura3; trp1-289; MAL2-8c; SUC2; XI-1-LoxP-KlLEU2-PaAroZ<-PTDH3-PTEF1->CaCatA; X-4-loxP-TRP1-ScTkl1<-PTDH3-PTEF1->KpAroY.D; pXII-4-LoxP-HphMXsyn-209bp_CYC1p_BenO_T1->yEGFP; pX-3-KanMXsyn-REV1p::BenM-MP17_D08; XII-5-KpAroY.B<-PTEF1-PTDH3->KpAroY.Ciso_C150W | N/A | Sc-78 | This study |

**Supplementary Figure S1. Graphical overview of p1 sequence after pMB11 insert integration.**

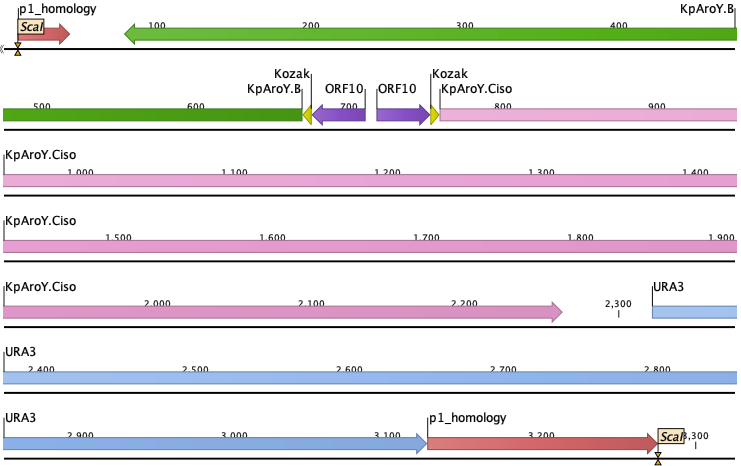

**Supplementary Figure S2. Graphical overview of wild-type genome integration cassette for reverse engineering with homology regions shown.**

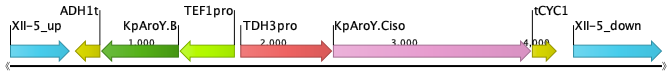

**Supplementary Figure S3. Graphical overview of identified mutations in sequenced p1 amplicons.**

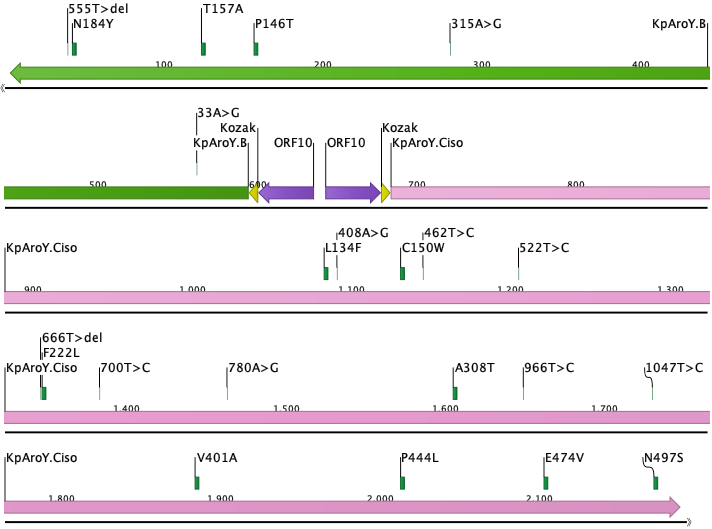

**Supplementary Information references**

DiCarlo, J.E., Norville, J.E., Mali, P., Rios, X., Aach, J., and Church, G.M. (2013) Genome engineering in Saccharomyces cerevisiae using CRISPR-Cas systems. *Nucleic Acids Res* **41**: 4336–4343.

Jessop-Fabre, M.M., Jakočiūnas, T., Stovicek, V., Dai, Z., Jensen, M.K., Keasling, J.D., and Borodina, I. (2016) EasyClone-MarkerFree: A vector toolkit for marker-less integration of genes into Saccharomyces cerevisiae via CRISPR-Cas9. *Biotechnol J* **11**: 1110–1117.

Ravikumar, A., Arzumanyan, G.A., Obadi, M.K.A., Javanpour, A.A., and Liu, C.C. (2018) Scalable, Continuous Evolution of Genes at Mutation Rates above Genomic Error Thresholds. *Cell* **175**: 1946–1957.e13.

Skjoedt, M.L., Snoek, T., Kildegaard, K.R., Arsovska, D., Eichenberger, M., Goedecke, T.J., et al. (2016) Engineering prokaryotic transcriptional activators as metabolite biosensors in yeast. *Nat Chem Biol* **12**: 951–958.

Zhong, Z., Wong, B.G., Ravikumar, A., Arzumanyan, G.A., Khalil, A.S., and Liu, C.C. (2020) Automated Continuous Evolution of Proteins in Vivo. *ACS Synth Biol* **9**: 1270–1276.
